# Short linear motif based interactions and dynamics of the ezrin, radixin, moesin and merlin FERM domains

**DOI:** 10.1101/2020.11.23.394106

**Authors:** Muhammad Ali, Alisa Khramushin, Vikash K Yadav, Ora Schueler-Furman, Ylva Ivarsson

## Abstract

The ERM (ezrin, radixin and moesin) family of proteins and the related protein merlin participate in signaling events at the cell cortex. The proteins share an N-terminal FERM (band Four-point-one (4.1) ERM) domain comprised of three subdomains (F1, F2, and F3) that hold multiple binding sites for short linear peptide motifs. By screening the FERM domains of the ERMs and merlin against a phage library that display peptides representing the intrinsically disordered regions of the human proteome we identified more than 220 FERM binding peptides. The majority of the peptides contained an apparent Yx[FILV] motif, but ligands with alternative motifs were also found. Interactions with thirteen peptides were validated using a fluorescence polarization assay, and interactions with seven full-length proteins were validated through pull-down experiments. We investigated the energy landscapes of interactions between the moesin FERM domain and representative set of ligands using Rosetta FlexPepDock computational peptide docking protocols, which provide a detailed molecular understanding of the binding of peptides with distinct motifs (YxV and E[Y/F]xDFYDF) to different sites on the F3 subdomain. A third motif (FY[D/E]L(4-5x)PLxxx[L/V]) was proposed to bind more diffusely. By combining competition and modeling experiments, we further uncovered interdependencies between different types of ligands. The study expands the motif-based interactomes of the ERMs and merlin, and suggests that the FERM domain acts as a switchable interaction hub where one class of ligands to the F3 subdomain allosterically regulates binding of other F3 ligands.

## INTRODUCTION

The ERMs (ezrin, radixin and moesin) are membrane-associated proteins that provide linkage between membrane and actin cytoskeleton [1, 2]. The proteins have important roles in controlling the localization of peripheral membrane proteins and the signaling from membrane receptors. The ERMs are closely related (**Fig. 1A**) and share an N-terminal FERM (F for 4.1 protein, E for ezrin, R for radixin and M for moesin) domain, followed by a region with ⟨α-helical propensity, and a C-terminal domain (CTD) that binds to F-actin (**Fig. 1B**). The FERM domain has a cloverleaf-like structure with three subdomains (F1, F2, and F3; **Fig. 1C**) [3]. The interdependent subdomains take distinct structures, with F1, F2 and F3 forming ubiquitin-like, acyl-CoA binding protein like, and phosphotyrosine binding (PTB) like folds, respectively. The protein merlin, encoded by the NF2 gene, is closely related to the ERMs (**Fig. 1A)**. Merlin is a well-known tumor suppressor protein and an upstream regulator of the Hippo pathway [4]. Compared to the ERMs, merlin lacks the C-terminal F-actin binding region (**Fig. 1B**) and has distinct tissue localization and function [5].

**Figure 1.**
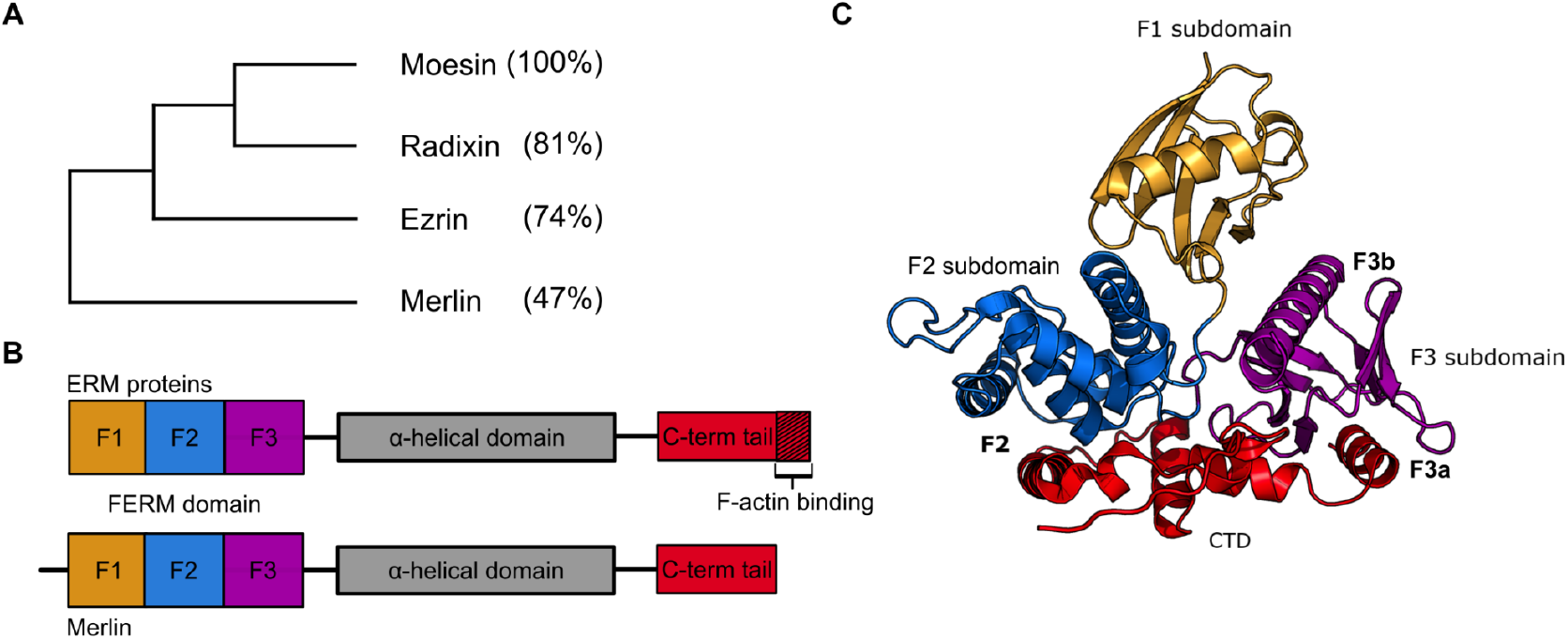
Overview of the ERM and merlin subfamily of the FERM domain containing proteins investigated. **(A)** Sequence identities between full-length proteins. (**B)** Modular architecture of the ERMs and merlin. (**C**) Structure of moesin in complex with its CTD. The different subdomains and the binding sites are indicated (PDB: 1EF1 [3]). Colors are according to the domain architecture shown in (B).

The FERM domain serves as a dynamic hub that binds to membrane phospholipids [6, 7] and short linear motif (SLiMs) containing binding partners, including intracellular regions of transmembrane receptors (**Table 1**). SLiMs are typically 3-10 amino acids long and commonly found in the intrinsically disordered regions of target proteins [8]. Distinct SLiM-binding sites have been found distributed over the different subdomains. Importantly, the F3 subdomain has two closely located but distinct binding pockets (F3a and F3b, **Fig. 1C**). Using the available complexes of peptide-bound FERM domains, ligands can be classified based on their binding pocket preferences (**Table 1**). Following this classification, the LLY motif from matrix the metalloproteinase-14 (MT1-MMP) belongs to class F1 [9], the -LxEI**-** containing peptide (where x = any amino acid) from the serine/threonine-protein kinases LATS1/2 belongs to class F2 [10], and the MDWxxxxx[L/I]Fxx[L/F]-coo-motif from the C-termini of the Na(+)/H(+) exchange regulatory cofactor NHE-RF1/2 (NHERF-1, also called EBP50) belongs to class F3a [11, 12]. Two different classes of ligands have been found to bind to the F3b pockets of moesin and radixin and are here defined as class F3b^1^ (Lx[I/L]N) [13] and class F3b^2^ (Yx[V/P]) (**Table 1**) [14, 15]. Notably, while both the ERMs and merlin FERM domains bind to LATS1/2 and NHERF-1/2 using their F2 and F3a pockets, they have distinct F3b pocket specificities [15, 16].

**Table 1.** Overview of ERM and merlin peptide ligands with known FERM binding sites based on crystal structures of complexes. Bold letters indicate key residues implicated in binding.

| FERM domain protein | Ligand | Peptide Sequence | Binding pocket | PDB code |
| --- | --- | --- | --- | --- |
| **Radixin** | MMP14 | RRLLYCQ | F1 | 3X23 [9] |
|  | NHERF-1 | **MDWSKKNELFSN**<br>**L** | F3a | 2D10 [12] |
|  | NHERF-2 | **MDWNRKREIFSN**<br>**F** | F3a | 2D11 [12] |
|  | CD43 | TG**ALT**LS | F3b<sup>1</sup> | 2EMS [59] |
|  | CD44 | KKKL**V**IN | F3b<sup>1</sup> | 2ZPY [13] |
|  | NEP | **MDITDIN** | F3b<sup>1</sup> | 2YVC [60] |
|  | ICAM2 | **RTGTYGV**LAA | F3b<sup>2</sup> | 1J19 [14] |
|  | PSGL-1 | KTHMYPVRNY | F3b<sup>2</sup> | 2EMT [61] |
| **Moesin** | CTD | - | F2, F3a | 1EF1 [3] |
|  | NHERF-1 | MDWSKKNELFSN<br>**L** | F3a | 1SGH [11] |
|  | CD44 | KKKL**V**IN | F3b<sup>1</sup> | 6TXS |
|  | CRB | ATRGT**Y**SPSA | F3b<sup>2</sup> | 4YL8 [15] |
| **Merlin** | CTD | - | F2, F3a | 3U8Z [62], 7EDR [63], 4ZRJ [10] |
|  | LATS1 | KALQEIRNS**LLPF** | F2 | 4ZRK [10] |
|  | LATS2 | KALREIRYS**LLPF** | F2 | 4ZRI [10] |
|  | DCAF1 | DIILSLN* | F3b | 3WA0 [64], 4P7I [16] |

\* A longer $\beta$ -hairpin motif was suggested by Li and co-authors [16].

The ERM proteins are regulated by autoinhibitory interactions between their C-terminal regions and their FERM domains (**Fig. 1C**) [17-21]. In the closed conformation, the C-terminal region of the proteins block both the F2 and the F3a sites of the FERM domains (**Fig. 1C**). Additional regulation may be provided by allosteric inhibition between different binding sites. Indeed, by comparing the unbound structure of radixin to a structure with its F3a pocket occupied by the EBP50 peptide it was found that peptide binding caused a conformational change that narrowed the F3b site [12]. Pre-incubation of radixin FERM with the EBP50 peptide was consistently shown to inhibit binding of class 3b ligands [12]. If and how class 3b ligands in turn affect EBP50 binding has to our knowledge not been described. In contrast, the F3b pocket and the F1 pocket appear to act independently of each other [9].

To better understand the function of the FERM domains we screened them against a peptide-phage library that tiles the intrinsically disordered regions of the human proteome [22]. We identified 222 ERM binding peptides and a smaller set of merlin ligands. We showed by modelling and mutational analysis that peptides with YxV motif bind to the F3b pocket, consistent with previous findings. Based on competition experiments and modelling we further propose a novel class F3a motif (E[Y/F]xDFYDF). Moreover, a third FY[D/E]L(4-5x)PLxxx[L/V]) motif was found. Through competition experiments and modelling we found that binding of F3a ligands block binding to other F3 sites. Taken together, the study contributes to an improved understanding of the interactions and dynamics of the FERM domains.

## RESULTS & DISCUSSION

### The ERM proteins bind Yx[FILV] containing peptides from cell adhesion related proteins and proteins involved in transcriptional regulation

To capture potential SLiM-based interactors, we used the FERM domains of the ERMs and merlin as baits in selections against a previously described peptide-phage library that display peptides representing intrinsically disordered regions of the human proteome (HD) [23]. Binding-enriched phage-pools were analyzed by next-generation sequencing (NGS). The sequences were translated into peptides and analyzed following an established protocol [24]. Peptides were assigned with confidence scores (0: no confidence; 1: low, 2-3: medium and 4: high) based on four metrics, namely the i) occurrence of peptides in replicate selections, ii) identification of motifs with overlapping peptides, iii) high NGS counts and iv) the presence of consensus motif. For a stringent analysis we focused on the medium/high confidence set of ligands (**Tables S1-S3**). Between 106 and 143 unique peptides were identified as high/medium ligands for each of the ERM FERM domains. The ligand sets overlapped, and the total number of unique high/medium confidence FERM domain binding peptides were 222 (**Fig. 2A**; **Tables S1-S3**). Consensus motifs were generated based on the binding enriched peptides using the SLiMFinder algorithm [25], which revealed that the data of the ERM FERM domains was dominated by ligands with an apparent Yx[FILV] motif (**Fig. 2B**).

**Figure 2.**
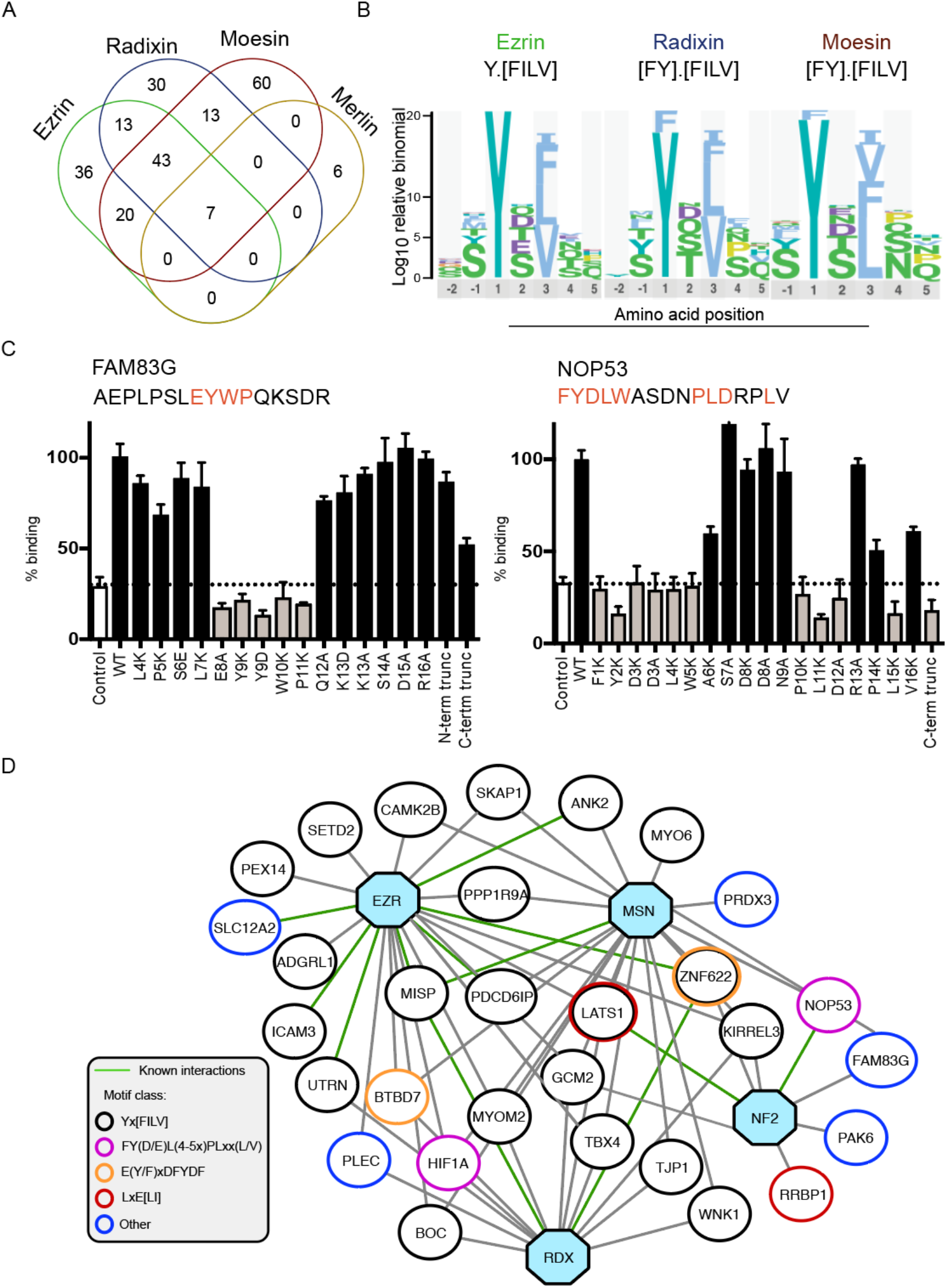
Proteomic peptide phage display results for the FERM domains of the ERMs and merlin. (**A**) Overlap of high/medium confidence peptide ligands identified through phage selections against the ERM proteins and merlin (Tables S1-3, S5). The merlin ligands include also low-confidence interactions validated through clonal phage ELISA (see Fig. S1). (**B**) Consensus binding motifs of the medium/high confidence ERM ligands generated by PepTools [24]. (**C**) Key amino acids in merlin binding FAM83G and NOP53 peptides identified through mutational analysis. Peptides were displayed on the p8 protein of the M13 phage. The effects of the mutations on binding were evaluated by clonal phage ELISA against immobilized GST-tagged merlin FERM domain. The binding was assessed by the ratio of the A_450_ values detected for the immobilized target protein (GST-tagged merlin) and the background (GST). The results were normalized to 100% binding of the respective wild-type peptide. As an extra negative control (indicated “control”) a clonal phage ELISA was performed for the same proteins using an M13 phage displaying no peptide. (**D**) Network of a select set of putative protein-protein interactions based on the proteomic peptide phage display results. Shown interactors of the ERMs and merlin include proteins that share biological functions with the baits that are unlikely to occur by chance based on GO term analysis (**Tables S1-S3, S5)**. The merlin interactors also include interactions validated through pull-down or FP binding experiments (**Table 2, Fig. 3**). The baits ezrin (EZN), radixin (RDX), moesin (MSN) and merlin (NF2) are indicated as blue octagonal. Interactions supported by results from other studies are indicated with green edges. The putative binding motifs of the binding peptides are indicated by the node color.

Through gene ontology (GO) term enrichment analysis of the combined set of ERM ligand using PepTools [24] we found an enrichment of peptides from protein associated with GO terms related to processes such as cell-cell adhesion, nervous system development and synapse organization (**Tables S1-S4**). There was also an enrichment of proteins involved in DNA binding. We visualized interactions with ligands that shared function with the ERMs (and/or) merlin that are unlikely to occur by chance based on the GO term analysis in a protein-protein interaction network (**Fig. 2D**). The network also includes interactions with novel SLiMs found in previously reported interactors of the ERMs, and interactions validated in the current study. Among the known interactions we note for example an interaction between ezrin and ankyrin-2 (ANK2) [26] previously found through proximity labeling mass spectrometry (AP-MS). Our results showed that ezrin can bind to ANK2 through three different binding sites (_2545-_EVS**Y**E**V**TPKTTDVSTP-_2560, 2950-_HTTSFHSSEV**Y**S**V**TIT-_2965_ and _3761-_PEESSLEYQQE**Y**F**V**TT-_3776_). Oligomeric ezrin may thus exhibit avidity effects upon binding to ANK2. The proteomic peptide-phage display analysis thus contributes with information on the binding motifs.

**Table 2.**
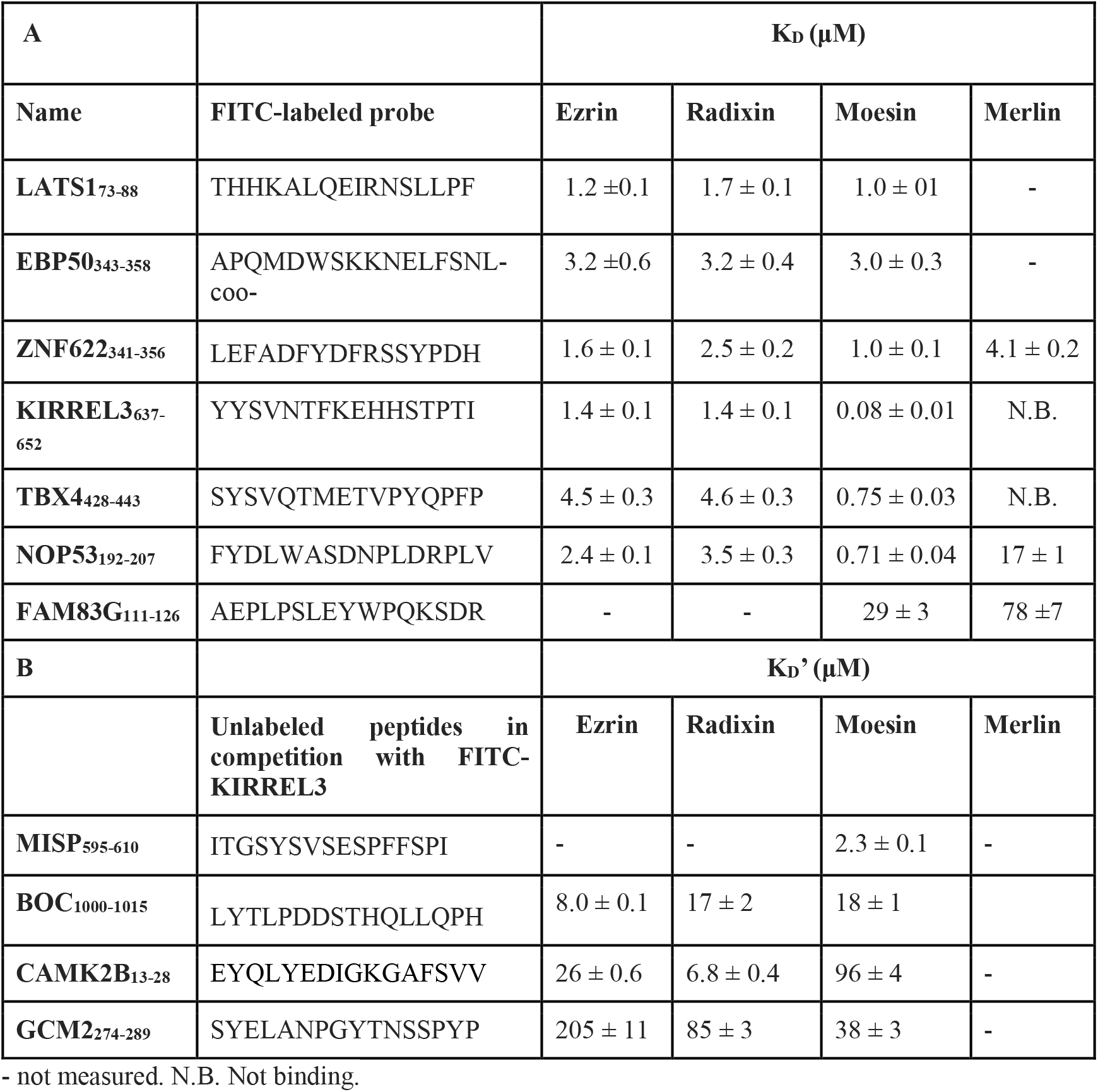
Affinities of the FERM domains of the ERMs and merlin for a representative set of ligands determined through direct binding FP experiments (A) or competition FP experiments (B).

| **A** |  | **K<sub>D</sub> (μM)** |  |  |  |
| --- | --- | --- | --- | --- | --- |
| **Name** | **FITC-labeled probe** | **Ezrin** | **Radixin** | **Moesin** | **Merlin** |
| **LATS1**<sub>73-88</sub> | THHKALQEIRNSLLPF | 1.2 ± 0.1 | 1.7 ± 0.1 | 1.0 ± 0.1 | - |
| **EBP50**<sub>343-358</sub> | APQMDWSKKNELFSNL-coo- | 3.2 ± 0.6 | 3.2 ± 0.4 | 3.0 ± 0.3 | - |
| **ZNF622**<sub>341-356</sub> | LEFADFYDFRSSYPDH | 1.6 ± 0.1 | 2.5 ± 0.2 | 1.0 ± 0.1 | 4.1 ± 0.2 |
| **KIRREL3**<sub>637-652</sub> | YYSVNTFKEHHSTPTI | 1.4 ± 0.1 | 1.4 ± 0.1 | 0.08 ± 0.01 | N.B. |
| **TBX4**<sub>428-443</sub> | SYSVQTMETVPYQFPF | 4.5 ± 0.3 | 4.6 ± 0.3 | 0.75 ± 0.03 | N.B. |
| **NOP53**<sub>192-207</sub> | FYDLWASDNPLDRPLV | 2.4 ± 0.1 | 3.5 ± 0.3 | 0.71 ± 0.04 | 17 ± 1 |
| **FAM83G**<sub>111-126</sub> | AEPLPSLEYWPQKSDR | - | - | 29 ± 3 | 78 ± 7 |
| **B** |  | **K<sub>D</sub>' (μM)** |  |  |  |
|  | **Unlabeled peptides in competition with FITC-KIRREL3** | **Ezrin** | **Radixin** | **Moesin** | **Merlin** |
| **MISP**<sub>595-610</sub> | ITGSYSVSESPFFSPI | - | - | 2.3 ± 0.1 | - |
| **BOC**<sub>1000-1015</sub> | LYTLPDDSTHQLLQPH | 8.0 ± 0.1 | 17 ± 2 | 18 ± 1 |  |
| **CAMK2B**<sub>13-28</sub> | EYQLYEDIGKGAFSVV | 26 ± 0.6 | 6.8 ± 0.4 | 96 ± 4 | - |
| **GCM2**<sub>274-289</sub> | SYELANPGYTNSSPYP | 205 ± 11 | 85 ± 3 | 38 ± 3 | - |

- not measured. N.B. Not binding.

### ProP-PD selection against merlin resulted in a small and diverse ligand set

For the merlin FERM domain, the ProP-PD analysis only resulted in 8 medium confidence peptides (**Table S5**). To gain more information on merlin ligands, we screened a set of low-confidence peptides for binding using clonal phage enzyme-linked immunosorbent assay (ELISA) (**Fig. S1**), which confirmed the interactions of additional 5 ligands, and thus increased the number of identified merlin ligands to 13. Of the total 13 putative merlin ligands, 7 overlapped with the ERM ligands. The merlin dataset contained two ligands with known or putative class F2 motifs, namely the know merlin binder LATS1 (_73-_THHKA**L**Q**EI**RNSLLPF-_88_) [10] and a novel merlin binding peptide from the ribosome-binding protein 1 (RRBP1; _1365-_RAATR**L**Q**EL**LKTTQEQ_-1380_). As the number of merlin peptides was not enough to establish consensus motif(s), we characterized the binding determinants of FAM83G_111-126_, the most enriched merlin ligand, and of the ribosome biogenesis protein NOP53 (NOP53_192-207_; previously shown to interact with merlin using its 181-479 region [27]) through a mutational analysis evaluated by clonal phage ELISA. A combination of lysine and alanine scanning was used for a clear readout in the relatively insensitive binding assay (**Fig. 2C**). For FAM83G_111-126_ the central -EYWP-stretch was necessary and sufficient for binding, as mutations of any of these residues abrogated binding and truncation of either the N-terminal region (AEPLP deletion) or the C-terminal region (KSDR deletion) only had minor effects. For NOP53_192-207_ the interaction required an extended FYDLWxxxxPLDxxL stretch. This was confirmed by the finding that a C-terminal truncation of the peptide (deletion of PLDRPLV) conferred loss of binding.

We enriched the merlin dataset for interactors of potential biological relevance, following the same principle as for the ERMs, and added these interactions to the network (**Fig. 2D**). The set of high confidence merlin ligands contained two that are from previously reported merlin interactors (LATS1 [10, 28] and NOP53 [27]) (**Fig. 2D**). The interaction between LATS1 and merlin contributes to the regulation of the Hippo pathway [28], and the nuclear interaction with NOP53 has been shown to confer growth inhibition of glioblastoma [27]. The most enriched merlin ligand FAM83G has been implicated in regulation of the bone morphogenetic proteins (BMP) pathway [29].

### The FERM domains bind to identified ligands with micromolar affinities

We determined the affinities (K_D_) of the FERM domains for a select set of ligands through a direct binding fluorescence polarization (FP) binding assay using a set of fluorescein (FITC)-labeled probe peptides, namely KIRREL3_637-652_ (KIRREL3: Kin of IRRE-like protein 3), TBX4_428-443_ (TBX4: T-box transcription factor TBX4), ZNF622_341-356_ (ZNF622: Zinc finger protein 622), NOP53_192-207_, and FAM83G_111-126_ (**Table 2A, Fig. 3 and Fig. S2**). The three first peptides were selected as variants of the Yx[FILV] motif (**Fig. 2B)**, and are from proteins that represent distinct biological functions: KIRREL3 is a synaptic adhesion protein [30], and the transcription factor TBX4 and the transcriptional activator ZNF622 are both involved in transcriptional regulation related to embryonic development [31, 32]. The latter two peptides (NOP53_192-207_, and FAM83G_111-126_) have distinct motifs as shown by the clonal phage analysis (**Fig. 2C**). As we in a next step aimed to explore the pocket specificity of the ligand we also included LATS1_73-88_ as a known F2 pocket ligand, and EBP50_343-358_ as a reference ligand for the F3a pocket (**Table 1**). The K_D_ values for interactions with the ERMs were in the low to mid micromolar range, except for the moesin-KIRREL3_637-652_ interaction, which was in a nanomolar range. For merlin, ZNF622_341-356_ was the highest affinity ligand, followed by NOP53_192-207_ and FAM83G_111-126_ The affinity for FAM83G_111-126_ was surprisingly low (78 µM) considering that it was the predominant ligand of merlin based on the phage display results. A plausible explanation for this may be that there is a bias for tryptophan containing peptides in phage display experiments [33]. Notably, the class F3b^2^ ligand KIRREL3_637-652_ did not bind merlin (**Table 2**), although being found in the merlin ProP-PD selection with low counts. This is in line with previous studies reporting that the two proteins have distinct F3b specificities (Table 1) [15, 16].

**Figure 3.**
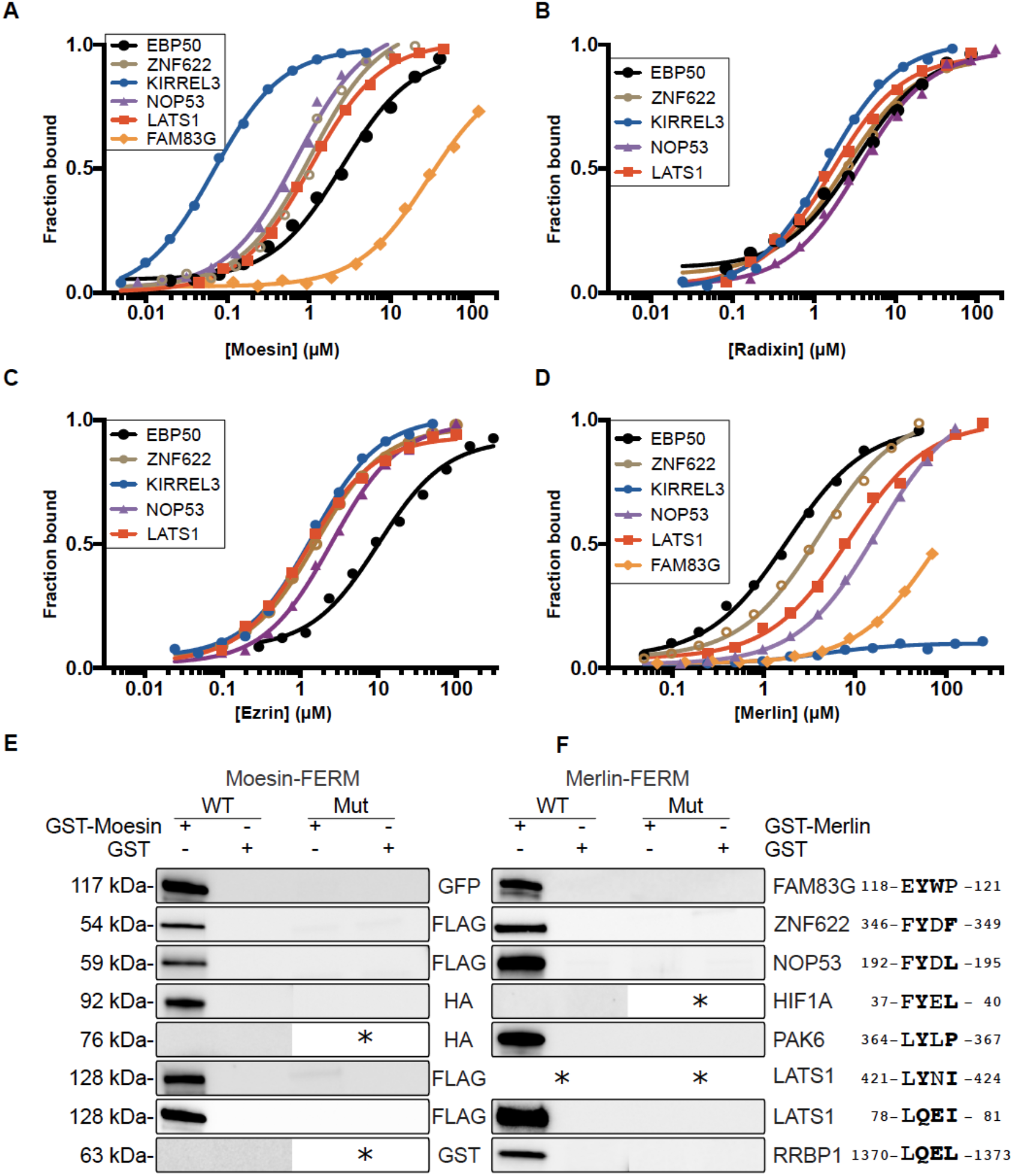
Determination of affinities through direct binding FP experiments and validation of interactions with full-length proteins through GST-pulldowns. **(A-D)** FITC-labeled probe peptides (5 nM) were titrated with increasing concentrations of (**A**) moesin, (**B**) ezrin, (**C**) radixin, and (**D**) merlin FERM domains. The results were fitted to a quadratic equation for 1:1 binding (see Table 2 for K_D_ values, n=3). The raw FP data is available in Fig. S2. (**E-F**) GST-pulldowns of wild-type and motif-mutant full-length proteins. GST-tagged moesin **(E)** or merlin **(F)** FERM domains were used to pull down wild-type or mutant proteins transiently expressed in HEK293 cells. Mutated amino acids are indicated to the right (bold residues were mutated to alanine). Note that LATS1 have two moesin binding motifs. Results shown are representative of at least two replicated experiments. * indicates not tested.

Interactions between the ERMs an additional set of four ligands were validated through a competitive FP assay where the probe peptide FITC-KIRREL3_637-652_ was outcompeted by increasing concentrations of unlabeled peptides (**Table 2B**). The observed transitions were fitted data to a sigmoidal dose versus response equation (**Fig. S3**), which generated IC50 values, from which apparent K_D_’ values were calculated. The analysis expanded the set of validated ERM ligands to the brother of CDO (BOC_1000-1015_), the calcium/calmodulin-dependent protein kinase type II subunit beta (CAMK2B_13-28_), the chorion-specific transcription factor GCMb (GCM2_274-289_) and the mitotic interactor and substrate of PLK1 (MISP_595-610_). Of these, MISP1 has previously been found to bind to the ERMs through high-throughput affinity-purification coupled to MS [34] and yeast-two-hybrid [35], but the binding region had remained unknown.

### Validation of interactions with full-length proteins and their binding motifs

The phage displayed peptides represent regions of human proteins and the selections thus identify putative interaction partners in the proteome. However, the binding interfaces may not be accessible in the context of the full-length proteins. We therefore validated interactions with seven full-length proteins through GST-pulldown experiments using moesin and merlin FERM domains as baits (**Fig. 3E-F**). Through these experiments, we confirmed that the moesin and/or merlin FERM domain interact with FAM83G, ZNF622, NOP53, HIF1A, PAK6, LATS1 and RRBP1. The target proteins have overlapping but distinct specificities for the FERM domains, such that an interaction with HIF1A was only confirmed for moesin, and the PAK6 and RRBP1 interactions were only confirmed for merlin. This is consistent with peptides from HIF1A being identified in selections against moesin, and PAK6 and RRBP1 being found in selections against merlin (**Tables S3, S5; Fig. 2D**). Through a mutational analysis we further confirmed that the identified binding motifs are crucial for the interactions with the full-length proteins. We validated the previously known LATS1 binding motif (_78-_LQEI_-81_) but also found a hitherto unknown second LATS1 FERM binding motif (_421-_LYNI_-424_), which appeared to reinforce the interaction with moesin (**Fig. 3E**).

### Exploring the structural basis of the interactions

We next used moesin as a model protein to explore which of the multiple binding sites the ligands bind to. Despite multiple attempts to co-crystallize moesin FERM with representative peptides we failed to obtain structures of complexes. We therefore turned to computational peptide docking using the Rosetta FlexPepDock modeling suite [36], using two main approaches: **i)** We applied FlexPepBind [37] to independently define the binding motif contained within the 16 amino acid long peptides. This protocol utilizes solved protein-peptide complexes onto which different peptide sequences can be threaded in order to differentiate between binders and non-binders, and in the case of a binder, to identify the bound peptide conformation. **ii)** Following the suggested motif definition, we applied a blind, global docking protocol - PIPER-FlexPepDock [38] to confirm the binding site and peptide conformation. In this approach we first select a predefined set of fragments that represents the peptide conformational ensemble. This set is then rigid-body docked to the receptor protein to sample all possible orientations of the peptide relative to the receptor. Finally the top-scoring complexes are refined using Rosetta FlexPepDock (See Methods).

For our simulations we used the available solved structures of moesin FERM domains, including the free moesin FERM domain (PDB ID 6TXQ [39]), the structure of moesin FERM domain with its own C-terminal domain (CTD) bound to the F3a and F2 sites (PDB ID: 1EF1 [3]), moesin with the crumbs CTD bound at the F3b site and in the cleft between the F3 and F1 lobes (PDB ID: 4YL8 [15]) and moesin with the CD44-derived peptide bound to the F3b site (PDB ID: 6TXS [40]).

### The YxV motif binds to the F3b binding site

We started the analysis with the YxV containing peptides from MISP_595-610_, KIRREL3_637-652_, and TBX4_428-443_ peptides (**Fig. 4**). These peptides have a class F3b^2^ like YxV motif similar to for example, the -**G**T**Y**G**V**-containing peptide from ICAM2 and the -**G**T**Y**S**P**-containing peptide of crumbs, which have been co-crystallized bound to the F3b site of radixin (PDB 1J19 [14]) and moesin (4YL8 [15]), respectively. Beyond the YxV motif, we noticed that MISP_595-610_ has a glycine at the p-2 position of the motif (-**G**S**Y**S**V**-), mimicking the glycine at the p-2 position of both the ICAM2 and the crumbs peptides. The glycine at the p-2 position allows tight packing of these peptides with moesin residue F250 (**Fig. 4B**) [15]. Starting from MISP_595-610_ we applied the FlexPepBind approach using two existing F3b bound structures (PDB codes 4YL8 and 6TXS) as templates. We threaded the sequence onto the peptides solved in these structures to identify the motif that could potentially bind at this site. The results converged for both simulations, identifying the IT<u>GS</u>**<u>Y</u>**S**V**S sequence as the best motif for the given binding site (**Fig. 4A,D)**. To reaffirm this binding mode, we globally docked the motif with PIPER-FlexPepDock. The top-scoring structure identified in this simulation hit the same site as the one identified by threading in a very similar binding conformation (**Fig. 4A**).

**Figure 4.**
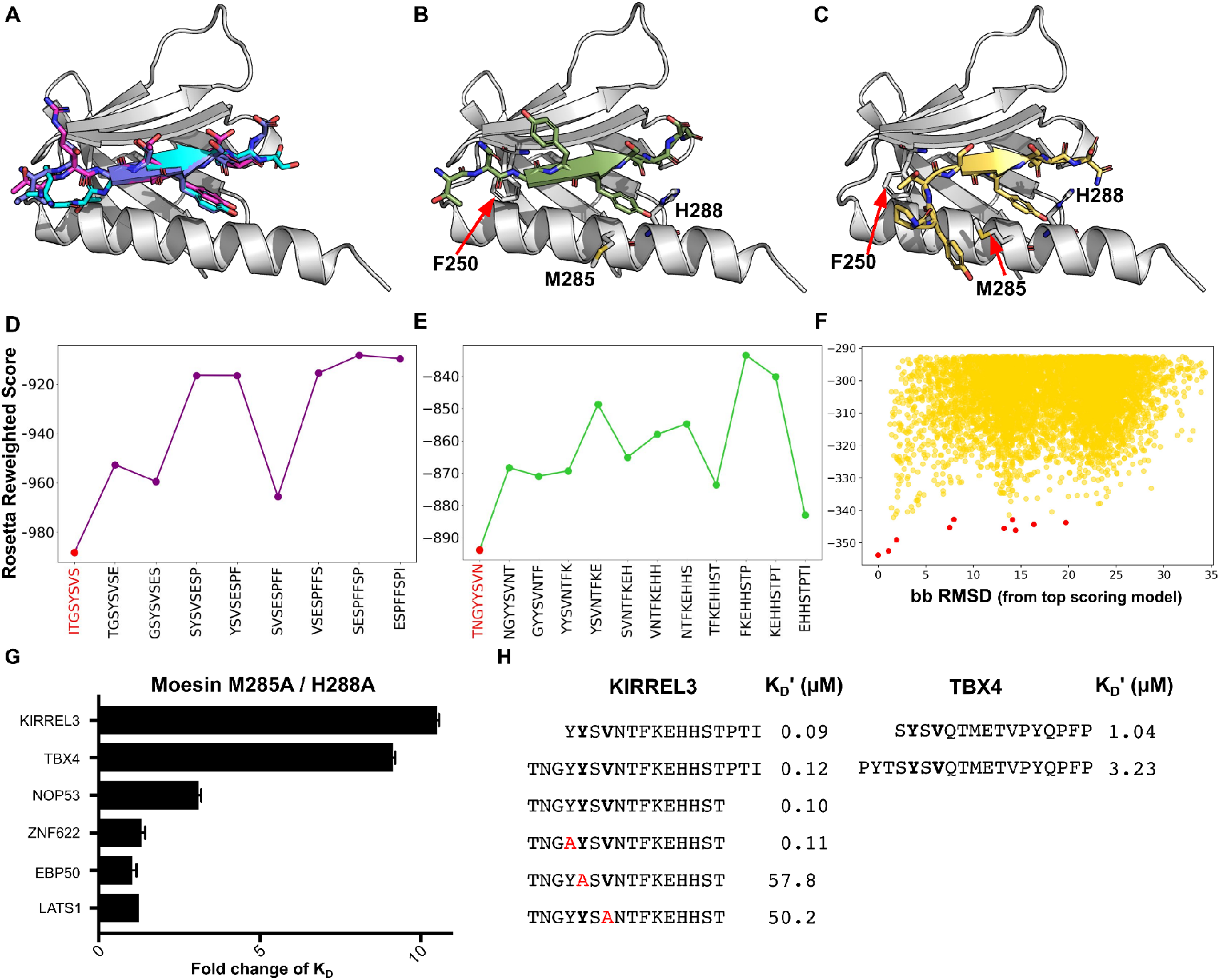
The MISP3, KIRREL3 and TBX4 top-scoring peptide models adopt similar conformations as the crumbs peptide in the F3b binding pocket. (**A**) Bound crumbs peptide (PDB 4YL8), MISP3 top-scoring models from FlexPepBind threading and PIPER-FlexPepDock global docking are shown in magenta, purple and cyan, respectively. (**B-C**) Top-scoring KIRREL3 peptide model from threading and TBX4 peptide model from global-docking are shown in green and yellow, respectively. The supposedly crucial residues that interact with the peptides are labeled. For each model, the corresponding energy landscapes are shown for threading and global docking simulations. (**D-E)** Threading binding energy landscapes of MISP and KIRREL, respectively. The identified motif is highlighted in red. (**F)** Global docking simulation binding energy landscape of TBX4, in which the lowest energy models converge towards the F3b site. Top 10 cluster representatives of each simulation are shown with red dots. (**G)** Fold change of the affinity of moesin FERM M285/H288A as compared to the wild-type (K_D_^M285/H288A^/K ^wt^) for six different peptides as determined by direct binding of FITC-labeled peptides (see Table 7 for complete data). (**H**) Competition FP experiment using variants of the KIRREL3 peptide (left) or the TBX4 peptide (right) for competition, and FITC-KIRREL3_637-652_ as probe.

In contrast, both threading and global docking of KIRREL3_637-652_ and TBX4_428-443_ failed to yield conclusive results. We therefore extended each peptide at the N-terminus to include the upstream residues, for KIRREL3 adding -TN<u>G-</u> to the -<u>S</u>**<u>Y</u>**S**V**-motif (TN**G**S**Y**SV), and for TBX4 adding -PYT-(PY<u>TS</u>**<u>Y</u>**S**V**S). The resulting KIRREL peptide featured glycine at position p-2 that matched the crumbs, MISP and ICAM2 sequences. We used the same strategy as for MISP_595-610_ to model the extended TBX4 and KIRREL3 peptides, namely FlexPepBind using the F3b bound template to identify the binding motifs, followed by global docking simulation of the best motif. For KIRREL3 the simulations converged in the same way as for MISP: the TN<u>GY</u>**<u>Y</u>**S**V**N motif identified by the FlexPepBind simulation was ranked best in the global simulation, adopting the same binding conformation as in threading **(Fig. 4B, E**). For the TBX4 peptide, threading still failed to identify any specific motif. However, global docking of the extended peptide (PY<u>TS</u>**<u>Y</u>**S**V**S) placed the peptide in a very similar position as the previous two peptides, with the N-terminus bulging out to accommodate the larger threonine residue **(Fig. 4C, F**).

To validate these docking models, we used computational alanine scanning to identify hotspot residues at the interface that would significantly affect the binding of the peptides, with a minor effect on the stability of the protein (**Table S6A**). Based on these calculations, we generated a mutant FERM domain with two mutations (M285A and H288A) and tested the effect on binding affinity. Consistent with the predictions, the mutations conferred a 9- and 10-fold loss of affinity for the TBX4 and KIRREL3 peptides, respectively, with no or minor effects on binding of the other probe peptides tested (LATS1, EBP50, ZNF622 and NOP53), supporting that only the YxV containing ligands bind to the F3b pocket (**Fig. 4G** and **Table S7**). We further confirmed the importance of the key residues of the YxV motif through a mutational analysis of the KIRREL3 peptide, and validated that the tyrosine at position p1 and valine at position p3, but not tyrosine at position p-1, are crucial for binding (**Fig. 4H**). The docking further suggested that the presence of a glycine at the p-2 position would improve the affinity. A glycine at p-2 was in fact present during the phage display of these peptides as the displayed peptides are flanked by a glycine-serine linker. We therefore experimentally tested how the N-terminal extensions of the KIRREL3 and TBX4 peptides affected the affinity of moesin **(Fig. 4H**). Notably, we found that the N-terminal extension of the KIRREL3 peptide did not change the affinity (K_D_ values for KIRREL3_637-652_ and KIRREL3_6334-652_ being 0.09 µM and 0.12 µM, respectively). In contrast, we noted a slightly reduced affinity for TBX4, which likely is due to a failure of its bulkier threonine at p-2 to pack against moesin F250, which forces the peptide to take a slightly different conformation, as suggested by the modelling (**Fig. 4C**).

### The E[Y/F]xDFYDF containing peptides ZNF622_341-356_ and BTBD7_940-950_ potentially bind to the F3a pocket

We next focused on the two peptides ZNF622_341-356_ and BTBD7_940-950_ that share an E[Y/F]xDFYDF stretch, and both bind to moesin with low micromolar affinities (**Table 2** and **Table S8**). To investigate how these peptides bind the FERM domain, we performed a global docking simulation on the full FERM domain, as well as on the F3 subdomain. Docking onto the full FERM domain positions the 10 top scoring models of the ZNF622 peptide at the F3a site, as well as in the cleft between the F3 and F1 domains (**Fig. S4**). In turn, docking the peptides onto the isolated F3 domain located the peptide in the F3a and F3b binding sites, strongly suggesting that this peptide binds to F3a. This is supported by the mutational analysis of the F3b site that showed that binding of ZNF622_341-356_ was unaffected (**Fig. 4G)**. The F3a bound peptide models did not converge to one single conformation, but rather adopted two distinct orientations: one similar to the part of the CTD bound at F3a, and another perpendicular to it (**Fig. 5A**). Notably, even though the backbone conformation of the peptide was different from the known binding mode at this site, the hydrophobic side-chains of the top-scoring models occupied the exact same sites as the CTD of moesin (**Fig. 5C**). The distinct preference between these two orientations is possibly determined by the flanking regions for CTD and EBP50 [11].

**Figure 5.**
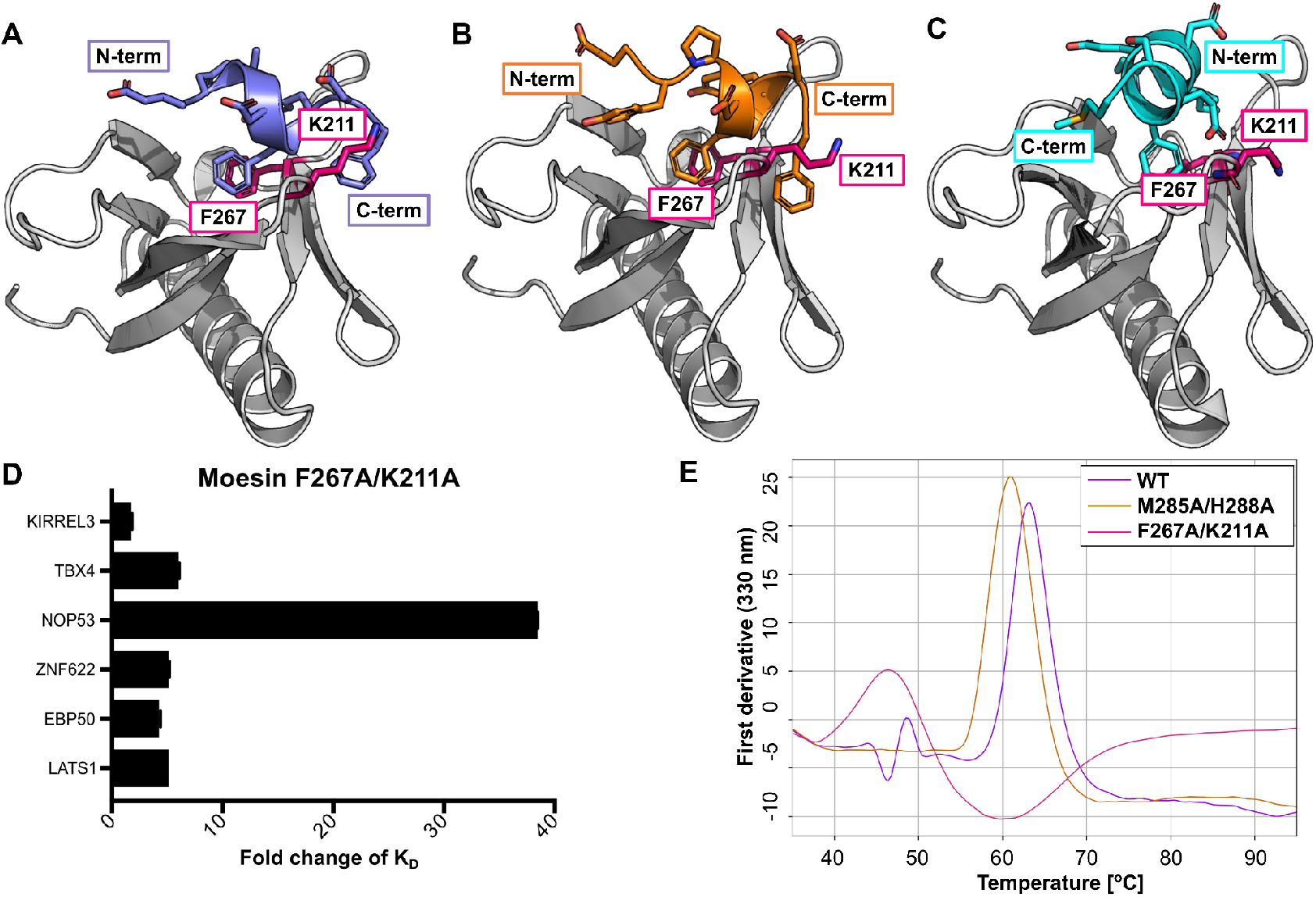
Structural modeling suggests that ZNF622 and BTBD7 bind the moesin FERM domain at F3a using the same binding pocket as moesin CTD and EBP50. **A-B**. Models of the interactions of ZNF622 (**A**) and BTBD7 (**B**) that both suggest a perpendicular conformation, both using aromatic side-chains to fill the same hydrophobic pocket. As expected, this distinct conformation is only identified by a global docking simulation but not by threading. (**C)** Fragment of moesin CTD bound at the same site (PDB code: 1EF1). (**D**) Mutational analysis of the proposed F3a binding site. Fold change of affinity for the moesin K211A/F267A mutant as compared to wild-type type moesin FERM domain (K_D_^M285/H288A^ /K ^wt^) for different FITC-labeled peptides. **(E)** Thermal unfolding profiles show that the F267A/K211A mutations (at F3a) destabilize the FERM domain, compared to WT and the M285A/H288A mutant at site F3b. Moesin mutants corresponding to potential F3a and F3b binding sites were analyzed for stability by thermal melting from 35 ºC to 95ºC.

Similarly, global docking simulation of BTBD7 onto the F3 subdomain of the moesin template placed most of the top-scoring models onto the F3a site, although no single defined binding conformation stood out. In addition, simulation on the full moesin template structure (after removal of the CTD) predicted the cleft between F3 and F1 binding lobes as the most probable binding site. Although the binding motif is very similar between the two peptides (BTBD7: EYP**DFYDF** vs. ZNF622: EFA**DFYDF**), PSIPRED [41] predicted the ZNF622 but not the BTBD7 peptide to form a helix, probably because of the presence of a proline upstream of the motif. Consequently, most of the starting fragments used for BTBD7 docking were in extended conformation. Assuming that both adopt a similar local conformation, we repeated our simulation with defined helical fragments. This simulation gave very similar results to those obtained for the ZNF622 peptide (**Fig 5B**).

To validate our docking models we identified three mutations that would perturb binding (K211A; I238A, F267A, **Table 6B**). Two of these mutations were tested in the format of a double mutant (moesin FERM K211A/F267A). The F267A mutation was chosen as it was predicted to have the largest effect on binding to both the ZNF622 and the BTBD7 peptides (although predicted to destabilize the protein, **Table S6B**) and the K211A mutation was added as it was proposed to reduce the affinity for the ZNF622 peptide. The double mutant exhibited a 5-fold loss in affinity for the ZNF622 peptide but also conferred similar loss of affinities for other probe peptides tested (e.g. the F2 binding FITC-LATS1_73-88_ and the F3 binding FITC-TBX4_428-443_), indicating a more general effect on the functional state of the protein (**Fig. 5D** and **Table S7**). Thermal shift analysis confirmed that the moesin FERM K211A/F267A mutant was destabilized (**Fig. 5E**), which suggests that the impaired stability is responsible for the general decrease in binding. Exactly how the E[Y/F]xDFYDF containing ligands bind to moesin thus remains to be validated by for example additional mutagenesis by the domain and the proposed binding motif, although the modelling converged on the F3a site. Of note, the K211A/F267A mutations had a significantly stronger, 30-fold effect on binding of the FITC-labeled NOP53_192-207_, consistent with its distinct binding mode, as described next.

### A diffuse binding pattern for the FY[D/E]L(4-5x)PLxxx[L/V]) containing peptides

Our mutational analysis of NOP53_192-207_ revealed that the peptide has an extended binding motif (FYDLWxxxxPLDxxL; **Fig. 2C**). Through inspection of the available peptides we found a similar motif in the HIF1A peptide (FY[D/E]L(4-5x)PLxxx[L/V]). Through FP competition experiments (described in the following section; **Table S8**) we found that the HIF1A_37-52_ peptide efficiently outcompeted FITC-NOP53_192-207_ but no other probe peptides, consistent with the two peptides binding in a similar way. Global docking of the NOP53 and HIF1A **FYDL**WASD and **FYEL**AHQL peptides respectively showed a diffuse distribution of binding modes that covered predominantly the F1-F3 cleft, and other interdomain sites (**Fig. S5**). The interaction with FITC-NOP53_192-207_ was affected the strongest by the destabilization conferred by the K211A/F267A mutations mentioned above (**Fig. 5E**). This might be explained by the proposed diffuse binding pattern. A binding strategy involving a number of weak binding sites which together result in low effective dissociation rate constants, and consequently, increased binding affinity, will be affected more severely by a general destabilization of the protein, reducing the affinity of the individual sites beyond the threshold needed for binding [42]

### Unveiling the interplay between different ligands

We next explored the interplay between different ligands using i) K_D_ measurements of FITC-labeled probe peptides in the presence of high concentrations of the competing ligands (**Fig. 6**) and ii) competitive FP experiments where the FITC-labeled probe peptide and the protein were kept constant and the probe peptides were outcompeted with increasing concentrations of unlabeled peptides (**Fig. 7**). We found that FITC-LATS1_73-88_ binding to the F2 site was largely unaffected by an excess of other ligands (EBP50_343-358_, KIRREL3_637-652_, TBX4_428-443_ or NOP53_192-207_) (**Fig. 6A,B**). However, there were some minor changes such that saturating concentrations of either EBP50_343-358_ or KIRREL3_637-652_ conferred a slightly increased affinity for LATS1_73-88_, while the opposite effect was observed for NOP53_192-207_. We further found that a high concentration of EBP50_343-358_ blocked binding of all other ligands tested. Surprisingly, we found that the opposite was not true, as binding of FITC-EBP50_343-358_ was largely unaffected by the presence of other ligands. The asymmetric results may partly be explained by an allosteric model where binding to the F3a site induces a conformational change of the protein that reduces the affinity for F3b ligands (**Fig. 6C**). The model is similar to the previously suggested allosteric inhibition of binding to the F3b site of radixin by EBP50 binding to the F3a site [12]. However, the lack of observable effect on the affinity of moesin for FITC-EBP50_343-35_ affinity by high concentrations of other ligands is not easily explained for a system in equilibrium. Nevertheless, the results consistently show that this was the case.

**Figure 6.**
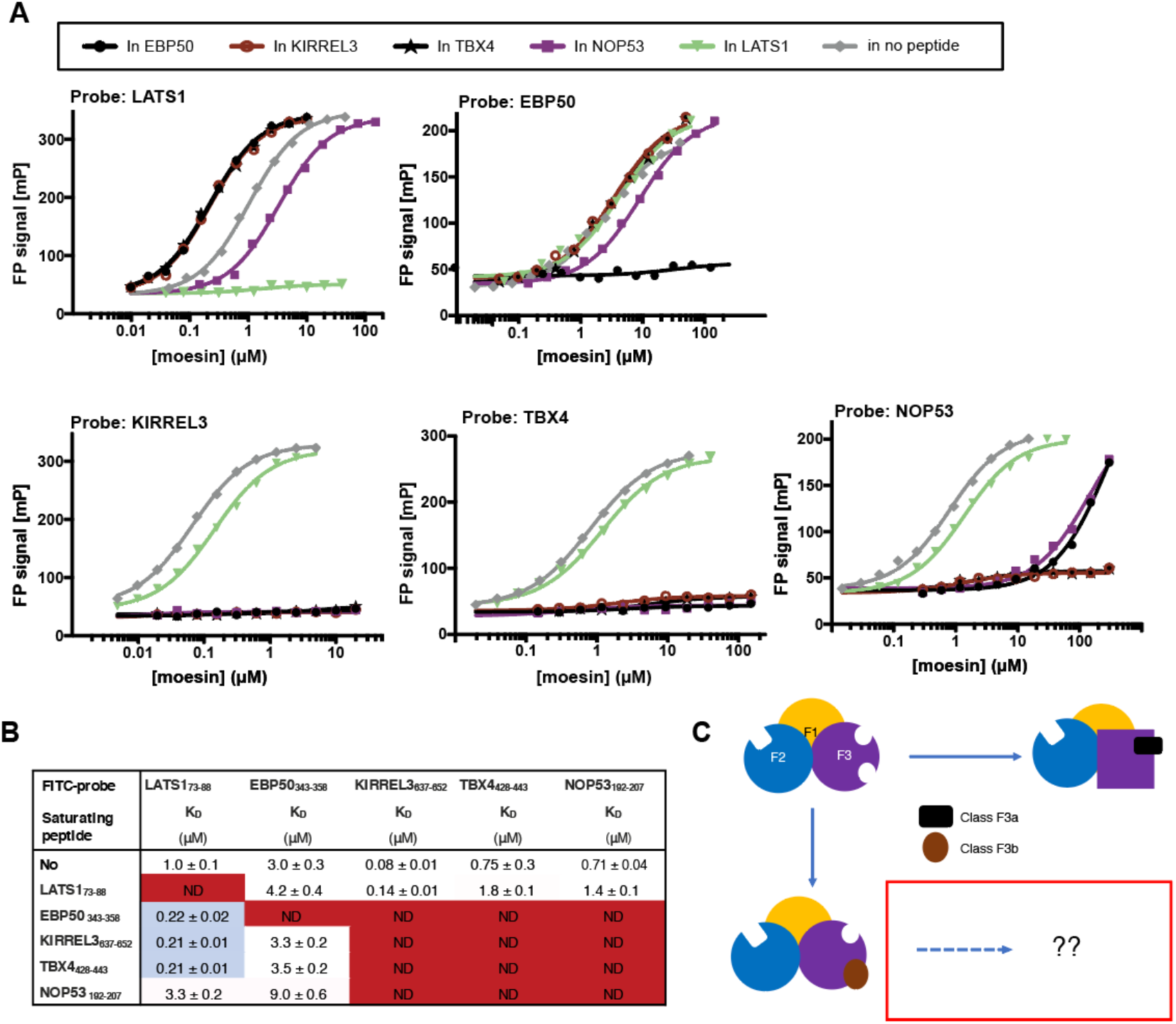
Asymmetric competition between EBP50_343-358_ and other ligands. (**A, B**) Direct binding of FITC-labeled probe peptides (5 nM; FITC-LATS1, FITC-EBP50_343-358_, FITC-KIRREL3_637-652_, FITC-TBX4_428-443_ and FITC-NOP53_192-207_) in the presence of saturating concentrations of competing ligands (80-150 µM; 50-100 x K_D_). A. Plots; B. Summary table (Red/blue squares indicate (close to) full competition/minor enhancement of binding). **(C**) Simplified schematic of the asymmetric effect observed between EBP50_343-358_ and other F3. Binding of EBP50_343-358_ to the F3a pocket hypothetically confers a conformational change of the domain (indicated as square) with low/no affinity for other F3 binding peptides. In contrast, binding of EBP50_343-358_ is unaffected by the presence of high concentration of F3b ligands.

**Figure 7.**
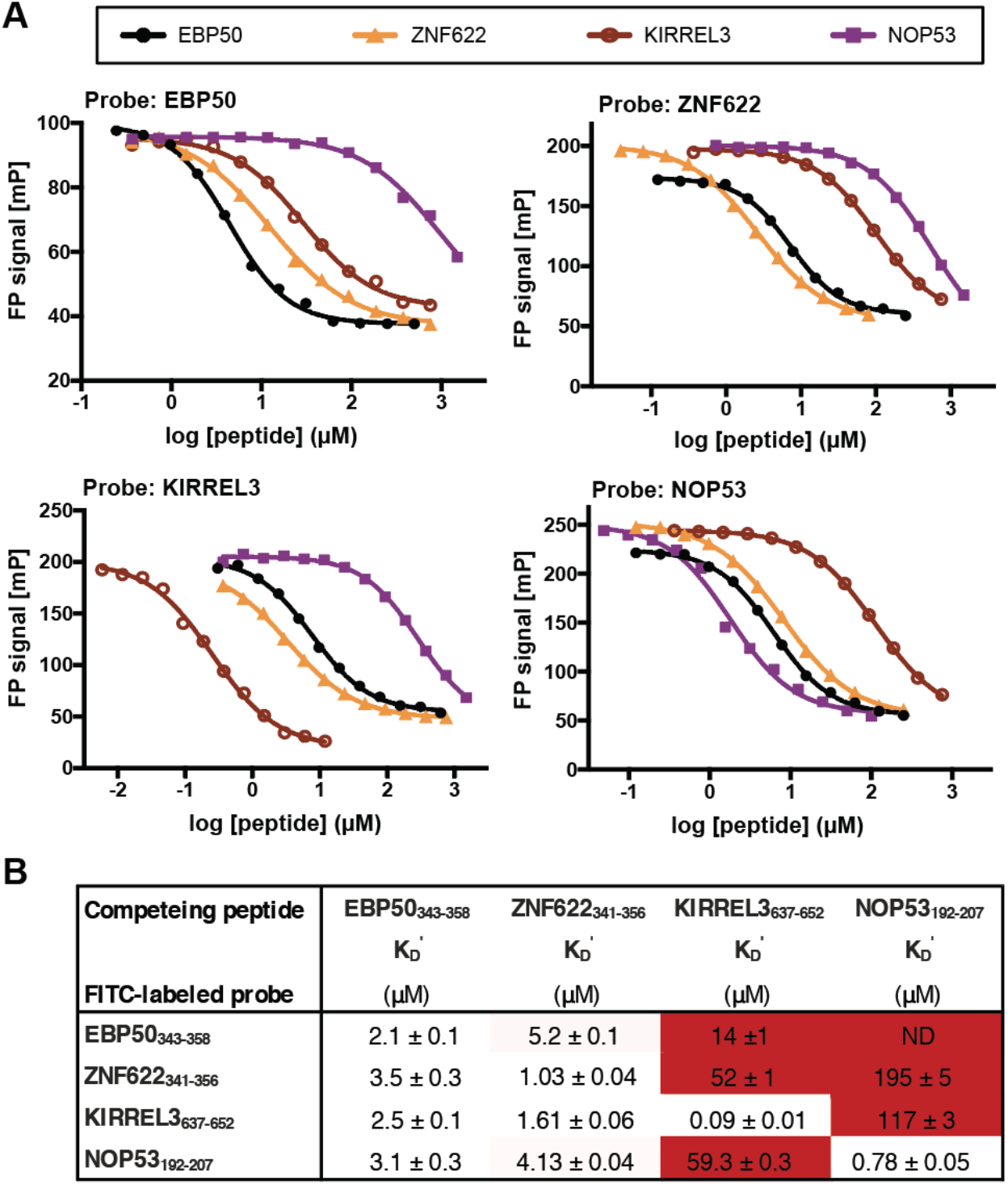
Competition between different ligands. (**A, B**) Competition between FITC-labeled probe peptides (5 nM; FITC-EBP50_343-358_, FITC-KIRREL3_637-652_, FITC-ZNF622_341-356_ or FITC.NOP53_192-207_ and unlabeled peptides (as indicated) for binding to moesin FERM domain (at a concentration of 1xK_D_ for indicated probe peptides). A. Plots; B. Summary Table (Red squares indicate that a large excess of competing peptide was needed to accomplish competition (K_D_’>>>K_D_)).

In further support of an asymmetric competition, we found through competitive FP experiments that unlabeled EBP50_343-358_ efficiently outcompeted moesin bound FITC-KIRREL3_637-652_ and FITC-NOP53_192-207_ with probe independent K_D_’ values (K_D_ values for direct binding was similar to the K_D_’ from competition experiments regardless of the probe peptides used) (**Fig. 7A,B**), In contrast, unlabeled KIRREL3_637-652_ and NOP53_192-207_ only could outcompete FITC-EBP50_343-358_ when used in high excess (**Fig. 7A,B)**. The FITC-EBP50_343-35_ binding was nevertheless readily reversible, as the FITC-EBP50_343-35_ probe was efficiently outcompeted with unlabeled EBP50_343-35_ as well as with the predicted F3a ligands ZNF622_341-356_ and BTBD7_940-950,_ (**Table S8**).

Taken together, there appear to be a complex mechanistic interplay between binding of different ligands that may involve a conformational change of the F3 domain upon binding to the F3a site, potentially paired with an additional step beyond the initial binding event that perturb the observed binding equilibrium.

### FERM F3 subdomain dynamics affects the affinity of the ligands

To test the suggested allosteric communication between the F3a and F3b sites, we checked how the use of the structures with different ligands occupying the distinct binding sites affect our docking simulations. We started with the ZNF622 peptide (predicted to bind at F3a), and checked whether it would still reach the F3a binding site on the receptor structure co-crystallized with CD44 at the F3b binding site (6TXS). This simulation identified F3a bound conformations among the top-ten models that are very similar to those from simulation on a corresponding F3a bound receptor structure (1EF1) (**Fig. (8A,B**). Additional models sampled the F3b binding site.

For the MISP peptide predicted to bind at F3b (**Fig. 4A**) we checked whether the peptide would still reach its binding site, when the F3a bound structure was used for the simulation (1EF1). This simulation was not able to localize MISP to the F3b site (the best peptide backbone RMSD among the top-ten cluster-representative structures is 5.9 Å away from the predicted binding conformation, **Fig. 8D**), despite successful identification of the F3b site when a bound receptor conformation was used (**Fig. 8E**). We were able to mimic the opening of F3b by relaxing a structure in presence of a superimposed F3b bound peptide (**Fig. S6**). Using that structure, a bound conformation was observed among the top-scoring structures at the F3b site (backbone RMSD of 2.2 Å from the top-scoring result of the bound simulation; **Fig. 8F**). This emphasizes the importance of opening the F3b binding site prior to docking, as mentioned previously.

**Figure 8.**
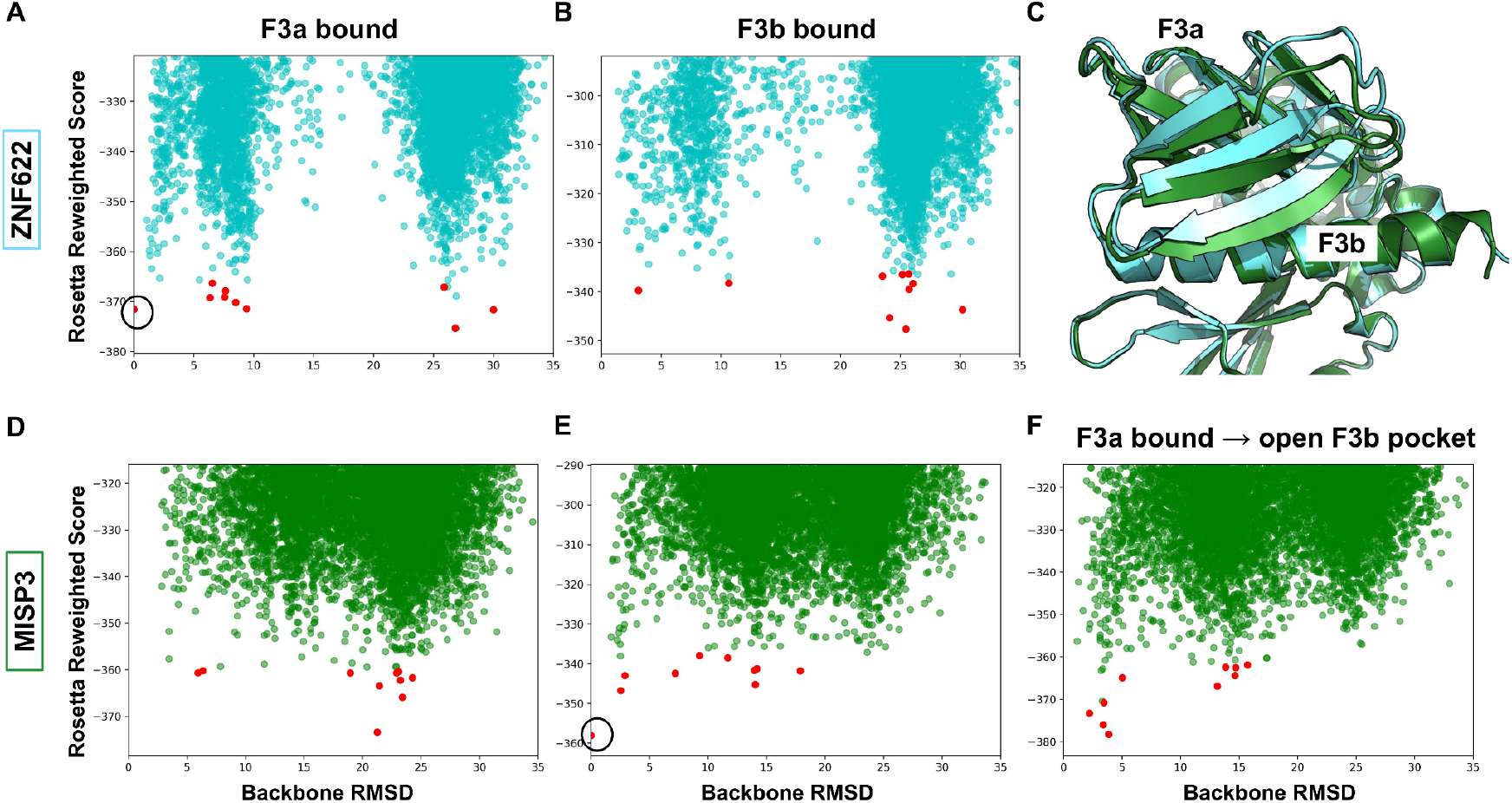
Energy landscapes of simulations using different receptor structures. (**A,B)**. Energy landscape sampled by the ZNF622 peptide: The F3a site is accessible both in F3a bound **(A)** and F3b bound **(B)** structures. **(C)** Comparison of the F3a bound (green) and F3b bound (cyan) structures shows conformational change depending on the occupied binding site. **(D,E)**: Energy landscapes demonstrate that the F3b site is inaccessible in the F3a bound FERM structure: Docking of the MISP peptide onto the F3b bound **(D)**, F3a bound **(E)**, and F3a bound relaxed structure with superimposed peptide at the F3b binding site (Fig. S6) (**F)** highlights the importance of conformational change for the identification of the F3b binding site. Cluster centers are shown with red dots. Only the F3 subdomain was used in the simulations. The highlighted circles in (A) and (E) show reference models used for RMSD calculations (structures shown in Figs. 5A and 4A, respectively).

In order to prove the generality of the influence of binding to the F3a site on the binding ability of the F3b, we performed docking simulations with ligands known to bind to these two pockets (i.e., with available solved structures: CD44 as a F3b ligand and CTD tail as a F3a ligand), using F3b-bound and F3a-bound structures. For CD44, the energy landscape shows no more top-scoring models within the F3b site in the F3a bound structure (see **Fig. S6**), due to F3b pocket closure (see **Fig. 8C**). In turn, no effect on the ability to model CTD binding to the F3a site was observed when using the F3b bound structure (**Fig. S7**). This data explains how the F3a binding peptides can outcompete all the other binders including those binding at a distinct pocket (i.e., F3b) by inducing a conformational change at the binding site, while binding to F3b will not affect binding to F3a.

### A distinct binding behavior for NOP53

As mentioned previously, NOP53_192-207_ shows a distinct binding pattern that is characterized by diffuse binding to a number of sites (**Fig. S5**), and most strongly affected by the destabilizing F267A/K211A mutation (**Fig. 5D,E**) (see above). Through direct FP binding experiments in presence of other ligands we observed a symmetric inhibition of binding between NOP53_192-207_ and F3b ligands TBX4_428-443_ and KIRREL3_637-652_ (**Fig. 6; Fig. 7**). The competitive FP experiment revealed that the apparent affinity of unlabeled KIRREL3_637-652_ for moesin as determined through competition against the FITC-NOP53_192-207_ probe was more than 600 times weaker than the affinity determined using the FITC-KIRREL3_637-652_ probe. Conversely, the apparent affinity of unlabeled of NOP53_192-207_ for moesin was more than 140 times weaker when using FITC-KIRREL3_637-652_ as probe instead of FITC-NOP53_192-207_. These results may be explained by the diffuse binding mechanism of NOP53_192-207_ that includes among others the F3b pocket, but also other sites. In turn, KIRREL3_637-652_ too could bind with weak affinity to alternative sites. This is well possible given the similarities of the binding motifs. Alternatively, the effects could be due to an induced fit mechanism where binding of one ligand reduces the affinity for a second site.

## CONCLUDING REMARKS

In this study we combined ProP-PD selections with cell based validations, biophysical affinity measurements and computational peptide docking to successfully expand the SLiM-based interactomes of the FERM domains of the ERMs and merlin and shed light on the complex interplay between different ligands. The ligands identified provide a rich resource for exploring the molecular function of the ERM proteins in cell-cell adhesion as well as in transcriptional regulation (**Table S4**). The latter might provide clues to the nuclear functions of the ERMs and merlin [43, 44]. The phage derived datasets contained representatives of at least four classes of FERM domain ligands (**Fig. 2D**). Using the information obtained from the existing structures, we built high-resolution models using Rosetta FlexPepDock-based methods and provided refined models of the interactions. Our models provide a structural basis for three different ways of interplay between binding partners: (1) Direct competition for the same binding site (e.g. KIRREL3 and TBX4 binding to F3b); (2) Allosteric intradomain communication between the F3a and the F3b sites (e.g. binding of EBP50 to F3a prevents of binding of KIRREL3 to F3b), and (3) diffuse binding (e.g. binding pattern of NOP53 and its mutual influence with KIRREL3).

Using the unconstrained Rosetta FastRelax protocol we were further able to model the conformational changes inside the domain caused by ligand binding to the F3a site. As the closed conformations of the ERMs and merlin involve the intramolecular binding of the C-terminal regions to the F2 and F3a sites this similarly leads to conformational changes that block also binding to the F3b site and potentially other sites. The full FERM domain proteins are consequently only binding available after activation by lipid binding and phosphorylation [7, 12, 45], and may in the open conformation be further regulated by the class F3a ligands. The potential interactions of the ligands with the ERMs in a cellular setting would thus be tightly regulated as most of the FERM domain ligands identified have class 3b^2^ motifs (**Fig. 2D**).

Not all interactions could be modeled as stable structures. For the NOP53 and HIF1A peptides, we were not able to identify strong binding preferences on the energy landscape (**Fig. S5**). This suggests that the peptides show a distinct binding pattern that is characterized by diffuse, weaker binding at various locations, which is strongly affected by the destabilization of the FERM domain (**Fig. 5D**). In turn, the observed diffuse binding pattern could also be due to the use of too short fragments in docking that do not represent well the biological interaction, especially since the mutational analysis of the NOP53 peptide revealed that residues downstream of the core motif are critical for binding merlin.

The present study contributes to the understanding of the ligand binding of one subgroup of FERM domains. The FERM family contains more than 50 additional members, and applying the here identified principles on the whole family may allow us to learn more about conservation, and variation of the binding pocket specificities as well as the allosteric communication within the FERM domain.

## MATERIALS AND METHODS

### Plasmids, cloning and mutagenesis

Synthetic genes encoding for human merlin FERM (22-312) and radixin FERM (5-295) domains were commercially synthesized (ThermoFisher) and cloned into pETM33 using NcoI and EcoRI restriction sites. pGEX4T Moesin-FERM was a gift from Vijaya Ramesh (Addgene plasmid # 1163 [46]). pGEX4T1 Ezrin FERM domain was a kind gift from Volker Gerke [47]. Human NOP53-HA-pcDNA was a kind gift from Ronit Sarid [48]. The NOP53 gene was PCR amplified and cloned into CMV10 using HindIII and EcoRI. RRBP1-GFP was kindly provided by Alexander F Palazzo [49]. HA-HIF1-alpha-pcDNA3 was a gift from William Kaelin (Addgene plasmid # 18949 [50]). pcDNA3 LATS1 was a gift from Erich Nigg (Addgene plasmid # 41156, [51]). LATS1 was PCR amplified and cloned into CMV10 using NotI and BamHI restriction sites. GFP-FAM83G-pcDNA was a kind gift from Gopal Sapkota [29]. HA-PAK6-pcDNA was kindly provided by Michael Lu [52]. ZNF622 was a gift from Hyunjung Ha [53]. All ligands for the pull down experiment were mutated at the binding site following the QuickChange mutagenesis protocol. The moesin FERM domain was mutated in two combinations of M285A/H288A and F267A/K211A. Mutations were made sequentially by following the QuickChange mutagenesis protocol. Sequences of all obtained, cloned and mutated constructs were confirmed using Sanger sequencing.

### Protein expression and purification

Expression constructs encoding the FERM domains of merlin, moesin, ezrin and radixin were transformed into *E. coli* BL21(DE3). For expression, 2xYT medium (1.6% tryptone, 1% yeast extract, 0.5% NaCl) was inoculated with a fresh overnight culture of the transformed cells and grown until the OD_600_ reached 0.8. Protein production was induced using 0.3 mM IPTG (isopropyl β-d-1-thiogalactopyranoside) by incubating cells overnight at 18ºC while shaking. Next day, cells were harvested by centrifuging at 7,000 xg for 10 minutes. For purification, the pellet was homogenized using PBS (137 mM NaCl, 2.7 mM KCl, 10 mM Na_2_HPO_4_, 1.8 mM KH_2_PO_4_, pH 7.4) containing lysozyme, DNase I, cOmplete™ protease inhibitor cocktail (Roche), 1% Triton X-100 and 2 mM ß-mercaptoethanol. Cells were lysed for 30 minutes while shaking at 4ºC followed by sonication for 5 cycles of 30 seconds. Cell debris was removed by centrifuging at 20,000 xg for 45 minutes and the supernatant containing His-GST-FERM domains was incubated with glutathione (GSH) sepharose resin (Cytiva) for 1 hour. Beads were collected and washed on column. The protein was eluted using 10 mM reduced GSH in PBS, pH 8.0 and dialyzed to PBS overnight using dialysis SnakeSkin dialysis tubing for subsequent use in phage selections and pull down experiments. Protein size and purity was analyzed by SDS-PAGE electrophoresis. The quality and stability of wild-type and mutant proteins were evaluated as described below.

For FP experiments of merlin FERM, the GST tag was removed using HRV3C protease by incubating the enzyme (100 units/10 mg protein) with protein overnight at 4ºC in the dialysis buffer (20 mM HEPES, pH 7.4, 150 mM NaCl, 0.05% Tween-20 and 3 mM DTT). The following day, the cleaved GST and the protease were removed using reverse Ni-IMAC and protein was concentrated to the working concentration. For FP experiments using moesin FERM, thrombin (Sigma) was used to cleave the GST-tag from moesin FERM domain on GSH sepharose beads (Cytiva) and incubated overnight at 4 ºC under gentle rotation. Next day, the beads were collected by gentle centrifugation while the supernatant containing thrombin and moesin FERM domain was passed through a HiTrap^®^ benzamidine column (Cytiva). The protein was then eluted using a high salt (0.7-1 M NaCl) concentration. The buffer was then exchanged using the PD-10 desalting column (Cytiva) into FP buffer (20 mM HEPES, pH 7.4, 150 mM NaCl, 0.05% Tween-20 and 3 mM DTT).

### Protein quality and stability

Purified moesin FERM domain was subjected to gel filtration using Superdex 200 (Cytiva) columns and the monomeric species was collected. During experiments, proteins were further subjected to various quality checks. W130i dynamic light scattering (DLS) (Avid Nano) was used before each set of moesin FP experiments to check if the protein was monomeric. Analytical gel filtration (Superdex 300GL (Cytiva)) also confirmed the monomeric state of moesin. The stability of WT and mutant moesin was analyzed by thermal unfolding using 5 µM of each protein on Tycho NT.6 (nanoTemper). Inflection temperatures (T_i_) were identified and relative stability was analyzed by comparing with WT moesin in the optimized buffer (20 mM HEPES, pH 7.4, 150 mM NaCl, 0.05% Tween-20 and 3 mM DTT), later used for FP measurements.

### ProP-PD selections

Phage selections were performed for four days using the HD library and method described in detail elsewhere [22, 54]. The library displays close to 500,000 human peptides on the p8 protein of the M13 phage. Three or more independent selections were performed for each bait proteins. For each replicate selection, 25 µg of target protein (GST-tagged FERM domains of merlin, moesin, ezrin and radixin) and GST-control were immobilized overnight to a 96-well maxisorp plate wells while shaking at 4 °C. Next day, the wells were blocked using 0.5% (bovine serum albumin) BSA in PBS for 1 hour. The naïve phage library containing 10^11^ colony forming units (CFU) was precipitated using 1/5^th^ volume of PEG/NaCl (20% PEG8000 and 0.4M NaCl) followed by centrifugation at 10,000 xg for 10 minutes and dissolved in 100 µl of PBS for each well. Control wells were washed four times using 200 µl PBS containing 0.05% Tween-20 (PBST) and incubated with phages for one hour at 4ºC with shaking to remove the non-specific binders. The target wells were washed as before and phage solution was transferred to them and incubated for 2 hours at 4°C. Unbound phages were removed by 4x washing with 200 µl PBST. The bound phages were eluted by adding 100 µl log phase *E*.*coli* OmniMax for 30 minutes at 37°C. M13KO7 helper phages (10^11^ pfu/ml) were added to each well and incubated again at 37 ºC for 45 minutes. Each hyper infected bacterial culture was transferred to 1 ml of 2xYT containing kanamycin (50 µg/ml), carbenicillin (100 µg/ml) and 0.3 mM IPTG and incubated overnight with shaking at 37°C. Following day, bacteria was pelleted by centrifugation at 5000 xg for 10 minutes and supernatant was used for phage precipitation as before. Phages were consequently dissolved in 1ml of PBS.

To determine the progress of the phage selections, a sandwich ELISA was performed. Briefly, 10 µg of target and control proteins were immobilized overnight in a 96-well maxisorp plate. Wells were blocked using 0.5% BSA. 100 µl of phage solution from each binding enriched phage pool was added to control and target wells and incubated for 1 hour at 4 ºC with shaking. Wells were washed 4 times using PBST and were incubated with anti-M13 coat HRP-conjugated antibody (1:5000) dilution for one hour. Unbound antibody was washed away as before and TMB substrate was added and allowed to develop the blue color. Reaction was stopped using 0.6 M H_2_SO_4_ and absorbance intensity was determined at 450 nm.

Peptide-coding regions of binding-enriched phage pools were amplified and barcoded using PCR. These DNA pools were analyzed using NGS on the Illumina platform (MiSeq) and results were demultiplexed and analyzed using a pipeline described elsewhere [24, 54]. The DNA sequences were translated to amino acid sequences, resulting in peptides associated with sequencing read counts. Confidence levels based on four metrics (occurrence of peptides in replicate selections, identification of motif containing regions with overlapping peptides, high NGS counts and the presence of consensus motif). Peptides that meet 2-4 of these criteria are considered of medium/high confidence, as benchmarked by Benz et al., [24].

### Fluorescence polarization assay

Synthetic peptides were obtained at 95% purity (GeneCust) and were dissolved in FP buffer (described earlier). To obtain direct binding saturation data, a dilution series of moesin FERM domain was prepared using the FP buffer and then an equal volume of 10 nM FITC-labelled peptide was added to each sample. After mixing, the FP signals were recorded using SpectraMax iD5 (Molecular Devices). Data was analyzed with GraphPad Prism version 7.0.0 for MacOS (GraphPad Software, San Diego, California USA). A quadratic equation for equilibrium binding [55] was used to fit the obtained data

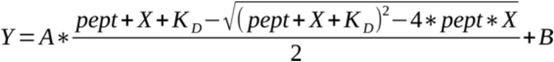

where *pept* indicates the probe peptide concentration (5 nM), *X* indicates the protein concentration, the constant *A* being the signal amplitude divided by probe peptide concentration, and B is the plateau value.

Direct binding experiments were performed and analyzed the same way in presence of saturating concentration of unlabeled peptides. In these experiments, the unlabeled peptide was used at a concentration approximately 100 times of the K_D_ values (80-150 µM concentrations). For the FP competition experiments, a pre-complex containing moesin FERM at 2x K_D_ concentration (for respective probe peptide) and 10 nM of FITC-labelled probe peptide was made in the FP buffer. 25 µl of this premix was added to 25 µl of titrated target peptide. FPs signals were determined as described early. Data was fitted to a sigmoidal dose-response (variable response; GraphPad Prism) model. The Hill coefficients were found to be around -1 for all experiments. All measurements were performed in at least triplicates.

### Cell culture and pull-down assays

HEK293 cells (Sigma:85120602) were cultured using DMEM (Gibco™) supplemented with 10% bovine FBS and NEAA (Gibco™) in a humid environment at 37ºC while maintaining 5% CO_2_. For pull-down experiments, the cells were transiently transfected with tagged-target proteins using Fugene^®^ HD (Promega) following manufacturer’s recommendations. Cells were allowed to grow and express the proteins for 48 hours post-transfection. Cells were washed with ice-cold washing buffer (PBS, Halt™ Protease Inhibitor Cocktail, pH 7.4) and then incubated with lysis buffer (50 mM Tris-HCl, pH 7.4, 150 mM NaCl, 10 mM sodium pyrophosphate, 10 mM sodium orthovanadate, 10 mM sodium fluoride, 1x cOmplete™ EDTA-free protease inhibitor tablet and 0.5% Nonidet P-40) for 30 minutes at 4ºC with gentle shaking. Cell debris was removed by centrifuging at 16,000 xg for 20 minutes at 4ºC and protein concentration was determined using BCA Protein Assay (Pierce™). Supernatant containing 0.5 mg of total protein was mixed with target FERM domain or GST (negative control) and one of the beads: GSH magnetic agarose beads (Pierce™), magnetic GFP-Trap® (Chromotek) and anti-FLAG® M2 magnetic beads (Sigma) according to manufacturer’s recommendations. This mixture was incubated overnight with end-over-end rotation at 4 ºC. Following day, the beads were collected and washed with lysis buffer 3 times. Samples were eluted using SDS-sample buffer by boiling at 95 ºC for 5 minutes.

Eluted samples were resolved by SDS-PAGE using 4-20% gradient gels and then transferred to the nitrocellulose membrane using the Trans-Blot Turbo transfer system (Biorad). Following the blocking for 1 hour in blocking buffer (5% milk in TBST), immunoblotting was performed using 1:2000 anti-FLAG (sigma), 1:2500 anti-GFP (ab6556), 1:2000 anti-HA (Sigma), 1:2500 anti-GST (Sigma) and 1:2000 anti-Myc (ab9106) antibodies followed by 1:5000 anti-rabbit HRP-conjugated secondary antibodies (GE) in blocking buffer. Membranes were exposed to Amersham ECL™ western blotting detection reagent (Cytiva) for a minute and signals were imaged using ChemiDoc™ Imaging system (Bio-Rad). Proteins with mutated putative binding sites were also immuno-precipitated and blotted similarly.

### Structure preparation for docking simulations

For peptide-protein interaction modeling two moesin crystal structures solved with different peptides were used: (1) F3b-bound (bound to CD44, PDB ID 6TXS), (2) F3a-bound (bound to CTD at the F3a site, PDB ID 1EF1). For the latter, we removed from 1EF1 the crystal contact peptide occupying the F3b site, which may bias the simulation towards the supposedly closed F3b site, and relaxed the structure using Rosetta FastRelax protocol [56] restraining heavy atoms to their native coordinates. Using this restrained “relax” protocol we attempted to remove any bias generated by the crystal contact.

### Global Blind Peptide Docking using PIPER-FlexPepDock

Global docking was performed using the PIPER-FlexPepDock protocol [38]. In brief, the peptide conformation is represented as an ensemble of fragments extracted from the PDB, based on sequence and (predicted) secondary structure using the Rosetta Fragment picker (with the vall2011 fragment library) [57]. These fragments are mutated to the target peptide sequence with the Rosetta fixed backbone design protocol [58]. 50 fragments are rigid-body docked onto the receptor protein using the PIPER rigid body docking program. The top 250 models for each fragment are then further refined using the Rosetta FlexPepDock protocol [36], including receptor backbone minimization, and top-scoring models are clustered. In this study all Rosetta simulations were performed using Rosetta version 2019.14. The protocol is freely available for noncommercial use as an online server: https://piperfpd.furmanlab.cs.huji.ac.il.

### Peptide threading with Rosetta FlexPepBind

The FlexPepBind protocol [38] uses a template structure of a protein-peptide interaction to thread a list of peptides each onto the template, and refines each peptide using FlexPepDock, and the top-scoring model is selected. In this study structural minimization only was used to refine the complexes, including both peptide and receptor backbone minimization. In some cases, where the template peptide was longer than the threaded peptide, the peptide sequences were threaded onto possible overlapping windows in the template.

### Modeling conformational changes with Rosetta FastRelax protocol

The Rosetta Relax protocol is used for full-atom refinement of protein structures. In this study the FastRelax protocol was applied with default parameters (no constraints were enforced) to open a pocket by superimposing the ligand to its binding site on an unbound structure. In the simulation 200 decoys were generated, and the top scoring model was taken for further analysis.

## Acknowledgements

This study was funded by grants from the Swedish research council (2016-04965 to YI), from the Carl Trygger foundation (YI, CTS14:209) and by the Israel Science Foundation, founded by the Israel Academy of Science and Humanities (grant number 717/2017 to OSF) and the US-Israel Binational Science Foundation 2015207 (to OSF). M was the recipient of a PhD fellowship from the Sven and Lilly Lawski foundation. Sequencing was performed by the SNP&SEQ Technology Platform in Uppsala. The facility is part of the National Genomic Infrastructure (NGI) Sweden and Science for Life Laboratory. The SNP&SEQ Platform is also supported by the Swedish Research Council and the Knut and Alice Wallenberg Foundation. The authors acknowledge the kind support of Norman Davey and Izabella Krystkowiak related to annotation of peptides and Leandro Simonetti for managing the NGS data.

## Author contributions

MA performed affinity determinations, molecular biology and cell-based experiments. MA and VKY performed phage experiments. AK performed the computational analysis, All authors analyzed results and conceived experiments. MA, AK, VKY, OSF and YI wrote the manuscript.

## Conflict of interest

The authors declare no competing interests.

## Notes

### Competing Interest Statement

The authors have declared no competing interest.

