## Supplemental Tables 1-8 for "Short linear motif based interactions and dynamics of the ezrin, radixin, moesin and merlin FERM domains"

### SUPPLEMENTAL TABLES S1-S8.

**Table S1.** Results of ProP-PD selections against Ezrin FERM domain.

| Confidence | uniprot_accession | gene | protein_name | designed_peptide | Total NGS |
|---|---|---|---|---|---|
| 3 | Q16602 | CALCRL | Calcitonin gene-relat | RSASYTVSTISDGPGY | 4140 |
| 2 | P55011 | SLC12A2 | Solute carrier family | AAARVELPGTAVPSVP | 2231 |
| 2 | Q8NEZ4 | KMT2C | Histone-lysine N-met | SYEVSSAPDVPSMGLV | 2221 |
| 2 | Q14938 | NFIX | Nuclear factor 1 X-ty | NFSLADLESPSYYNIN | 1082 |
| 2 | Q2LD37 | KIAA1109 | Transmembrane prot | EWTLDQPVSQTRTTAI | 643 |
| 3 | Q8WZ42 | TTN | Titin | VYSFEVQNPVGKDSCT | 524 |
| 3 | P49005 | POLD2 | DNA polymerase delt | FDPTNYTLPQQPLHPC | 435 |
| 2 | Q96BD5 | PHF21A | PHD finger protein 2: | TFTFPAPVQPVSLPSP | 433 |
| 3 | O60381 | HBP1 | HMG box-containing | QSDPTQSGMYQLSSDV | 388 |
| 3 | Q96BF6 | NACC2 | Nucleus accumbens-: | SYAVQEKPEPVPLESR | 386 |
| 2 | O60732 | MAGEC1 | Melanoma-associate | PVSSFFSYTLASLLQS | 375 |
| 2 | Q01484 | ANK2 | Ankyrin-2 | PEESSLEYQQEYFVTT | 368 |
| 2 | Q9HC84 | MUC5B | Mucin-5B | TWTVPAQTTTPMSTMS | 362 |
| 3 | Q9P203 | BTBD7 | BTB/POZ domain-coı | QEYPDFYDFSNAACRP | 343 |
| 3 | Q8IYH5 | ZZZ3 | ZZ-type zinc finger-c | DDDPDVYYFESDHVAL | 305 |
| 3 | Q6S9Z5 | ZNF474 | Zinc finger protein 47 | SSSGSYSLQATNEAAF | 234 |
| 2 | Q9Y5G0 | PCDHGB5 | Protocadherin gamm | QFTLQHVPDYRQNVYI | 209 |
| 2 | Q14653 | IRF3 | Interferon regulatory | IYEFVNSGVGDFSQPD | 206 |
| 2 | Q6PJT7 | ZC3H14 | Zinc finger CCCH dom | YYDMESMVHADTRSFI | 193 |
| 3 | Q8IVT2 | MISP | Mitotic interactor an | ITGSYSVSESPFFSPI | 168 |
| 2 | P01100 | FOS | Proto-oncogene c-Fo: | AFTLPLLNDPEPKPSV | 154 |
| 3 | Q9H9V9 | JMJD4 | JmjC domain-contain | YDVTSPALCDTHLHPR | 152 |
| 3 | P57082 | TBX4 | T-box transcription fa | SYSVQTMETVPYQPFP | 151 |
| 3 | Q9HBZ2 | ARNT2 | Aryl hydrocarbon rec | SSYDLSQVPVPNLPAG | 145 |
| 2 | Q86UR5 | RIMS1 | Regulating synaptic r | MYTLEHNDGSQSDTAV | 140 |
| 3 | Q8WYJ6 | SEPT1 | Septin-1 | LKEEEIHIYQFPECDS | 121 |
| 3 | O75603 | GCM2 | Chorion-specific tran: | SYELANPGYTNSSPYP | 121 |
| 2 | P22105 | TNXB | Tenascin-X {ECO:000 | TGSSPDSLSLSWTVPQ | 119 |
| 2 | Q9Y4K1 | CRYBG1 | Beta/gamma crystall | SFVLPVESTQDVSSQV | 118 |
| 3 | Q8IZM8 | ZNF654 | Zinc finger protein 65 | DYDLNQETSVIHKING | 112 |
| 2 | Q6PCD5 | RFWD3 | E3 ubiquitin-protein l | YFQVSRTQPDLPATTY | 112 |
| 3 | Q6NZI2 | CAVIN1 | Caveolae-associated | MEDPTLYIVERPLPGY | 110 |
| 3 | Q969S3 | ZNF622 | Zinc finger protein 62 | LEFADFYDFRSSYPDH | 109 |
| 3 | P20930 | FLG | Filaggrin | LYQVSTHEQSESAHGR | 101 |
| 2 | Q9Y264 | ANGPT4 | Angiopoietin-4 | SYTFLLPKSEPCPPGP | 100 |
| 3 | Q71RC2 | LARP4 | La-related protein 4 | SYQIYDVSGESNSAVS | 99 |
| 3 | Q01484 | ANK2 | Ankyrin-2 | EVSYEVTPKTTDVSTP | 95 |
| 2 | Q9UI33 | SCN11A | Sodium channel prot | QRITQPEPEQQAYELH | 86 |
| 3 | Q8IZU9 | KIRREL3 | Kin of IRRE-like prote | YYSVNTFKEHHSTPTI | 85 |
| 2 | P58107 | EPPK1 | Epiplakin {ECO:0000: | FFDPNTHENLTYVQLL | 85 |
| 2 | P46939 | UTRN | Utrophin | TVTEVDMDLDSYQIAL | 84 |
| 3 | Q13469 | NFATC2 | Nuclear factor of act | PLEWPLSSQSGSYELR | 82 |
| 3 | Q8IUX7 | AEBP1 | Adipocyte enhancer-l | FPFTTVETYTVNFGDF | 78 |
| 2 | Q7Z570 | ZNF804A | Zinc finger protein 8( | YTLIHSEENTKDATTV | 72 |
| 3 | P52292 | KPNA2 | Importin subunit alpl | ETTSEGYTFQVQDGAP | 69 |
| 2 | P35398 | RORA | Nuclear receptor ROI | TYNISANGLTELHDDL | 63 |
| 3 | A7KAX9 | ARHGAP3 | Rho GTPase-activatir | DYHVTQLQPYFENGRV | 58 |
| 3 | Q6EBC2 | IL31 | Interleukin-31 | LVSQNYTLPCLSPDAQ | 57 |
| 2 | O94766 | B3GAT3 | Galactosylgalactosyl | QPPEPEALPTIYVVTP | 57 |
| 2 | Q9HBV1 | POPDC3 | Popeye domain-conta | FYQMSTPEIRRSPLTQ | 56 |
| 2 | Q6ZVL6 | KIAA1549 | UPF0606 protein KIA | YYDFPAVETSKGLTER | 54 |
| 2 | Q7Z333 | SETX | Probable helicase sei | AYELSQRSLDYVAQLR | 50 |
| 2 | Q16665 | HIF1A | Hypoxia-inducible fac | FYELAHQLPLPHNVSS | 50 |
| 2 | Q9NRA1 | PDGFC | Platelet-derived grow | KYDFVEVEEPSDGTIL | 48 |
| 3 | Q14966 | ZNF638 | Zinc finger protein 63 | ILQLDSPESAQSMYSF | 45 |
| 2 | Q9BXT5 | TEX15 | Testis-expressed pro | TYQGITSYEVQPSPSG | 45 |
| 3 | Q96L42 | KCNH8 | Potassium voltage-ga | DYSPSHYQVVQEGHLQ | 42 |
| 3 | Q02505 | MUC3A | Mucin-3A {ECO:0000 | TSSITSTYTVTSMTTT | 42 |
| 2 | A8K2U0 | A2ML1 | Alpha-2-macroglobul | KYSMVELQDPNSNRIA | 41 |
| 2 | Q9Y6N5 | SQOR | Sulfide:quinone oxido | SYEMLHVTPPMSPPDV | 40 |
| 3 | Q99728 | BARD1 | BRCA1-associated RI | IKKDASAQQDSYEFVS | 39 |
| 2 | Q70E73 | RAPH1 | Ras-associated and p | SYSLDDVTAQLEQASL | 36 |
| 2 | Q8N0W7 | FMR1NB | Fragile X mental reta | TYELAATESNPESSHP | 36 |
| 3 | Q96FA7 | ZBED6CL | ZBED6 C-terminal-lik | VSEYSLNSPSPLQSPR | 35 |
| 2 | Q9BWV1 | BOC | Brother of CDO | LYTLPDDSTHQLLQPH | 35 |
| 2 | P60606 | CTXN1 | Cortexin-1 | MSATWTLSPEPLPPST | 35 |
| 2 | O95835 | LATS1 | Serine/threonine-pro | THHKALQEIRNSLLPF | 35 |
| 3 | Q8NB66 | UNC13C | Protein unc-13 homo | CHSDLQDDSESYDLTQ | 33 |
| 3 | Q8N895 | ZNF366 | Zinc finger protein 36 | CYEVEPYSPGLAPQSQ | 33 |
| 3 | Q9Y250 | LZTS1 | Leucine zipper putati | LPTHSTSSSYQLDPLV | 33 |
| 3 | P54296 | MYOM2 | Myomesin-2 | TYSVSVSDTDGVSSSF | 33 |
| 2 | O75381 | PEX14 | Peroxisomal membra | TYHLLGPQEEGEGVVD | 32 |
| 3 | Q5H9K5 | ZMAT1 | Zinc finger matrin-ty | SLNQQENNSGSYSVES | 31 |
| 3 | Q7Z6R9 | TFAP2D | Transcription factor A | HHQSFHYEFQHSHPAV | 29 |
| 3 | Q9UQR1 | ZNF148 | Zinc finger protein 14 | KDEYMVAEYAVEMPHS | 29 |
| 3 | Q9P2H0 | CEP126 | Centrosomal protein | SSLSNVTSNYDFVGQH | 29 |
| 2 | O94910 | ADGRL1 | Adhesion G protein-c | DTLPLNGNFNNSYSLR | 27 |
| 3 | Q8WVF1 | OSCP1 | Protein OSCP1 | GTNMYSVNQPVETHVS | 26 |
| 2 | Q9Y5P4 | COL4A3BI | Collagen type IV alph | SQVEEMVQNHMTYSLQ | 25 |
| 2 | Q15149 | PLEC | Plectin | FFDPNTHENLTYRQLL | 24 |
| 3 | Q86UW6 | N4BP2 | NEDD4-binding prote | PDDQVYSFLPSQDVNS | 21 |
| 3 | P32942 | ICAM3 | Intercellular adhesio | SGSYHVREESTYLPLT | 21 |
| 2 | O15259 | NPHP1 | Nephrocystin-1 | SEQTYDFLGEMRKNAV | 21 |
| 2 | Q99543 | DNAJC2 | DnaJ homolog subfar | FTPWTTEEQKLLEQAL | 20 |
| 4 | P24928 | POLR2A | DNA-directed RNA po | SYSPTSPSYSPTSPSY | 19 |
| 3 | Q01484 | ANK2 | Ankyrin-2 | HTTSFHSSEVYSVTIT | 18 |
| 2 | P29074 | PTPN4 | Tyrosine-protein pho: | NLSGYLSDYSFIPNQP | 17 |
| 2 | O60704 | TPST2 | Protein-tyrosine sulfo | YFQVNQNSTSSHLGSS | 17 |
| 2 | Q92797 | SYMPK | Symplekin | SFTPHQQAHIPNSIMT | 16 |
| 2 | Q02962 | PAX2 | Paired box protein Pa | SNDPVGSYSINGILGI | 16 |
| 2 | Q99569 | PKP4 | Plakophilin-4 | QYRVQECNYNRLQHAV | 15 |
| 2 | P78524 | ST5 | Suppression of tumo | SYQFPKLDRPTKQMRE | 15 |
| 2 | Q9Y5V3 | MAGED1 | Melanoma-associate | TYNFSQSLNANDLANS | 15 |
| 2 | Q15032 | R3HDM1 | R3H domain-containi | YYSVIPPGQQNNLSSS | 15 |
| 3 | Q9NRJ4 | TULP4 | Tubby-related protei | LASQSSYSLLSPPDSA | 14 |
| 2 | Q9ULJ8 | PPP1R9A | Neurabin-1 | NEYSVTGHYPLNLPSV | 14 |
| 2 | O75427 | LRCH4 | Leucine-rich repeat a | SYDLSDITQADLSRNR | 14 |
| 2 | Q9UPW0 | FOXJ3 | Forkhead box protein | SYTPVTSHPESVSQSL | 14 |
| 2 | Q9Y581 | INSL6 | Insulin-like peptide IN | YSPYQFESPQTASPAR | 13 |
| 2 | Q8N2C7 | UNC80 | Protein unc-80 homo | NFTLPSPVLGMPSVPM | 12 |
| 2 | Q9NW82 | WDR70 | WD repeat-containin | TLTQDYIITPHALPMF | 12 |
| 2 | Q9UNW8 | GPR132 | Probable G-protein c | LQSPVALADHYTFSRP | 11 |
| 2 | Q6ZW49 | PAXIP1 | PAX-interacting prote | QSSERSEMIATWSPAV | 11 |
| 2 | P02675 | FGB | Fibrinogen beta chair | TSEMYLIQPDSSVKPY | 11 |
| 2 | Q3SXY7 | LRIT3 | Leucine-rich repeat, i | YTVIQESPEEGVRWSI | 11 |
| 2 | P02751 | FN1 | Fibronectin | PDVRSYTITGLQPGTD | 10 |
| 2 | O75676 | RPS6KA4 | Ribosomal protein S6 | FFQQYELDLREPALGQ | 9 |
| 2 | Q8IZU9 | KIRREL3 | Kin of IRRE-like prote | QNLKDPTNGYYSVNTF | 9 |
| 2 | Q9Y2H5 | PLEKHA6 | Pleckstrin homology | HYDVDINKELSTPDKV | 8 |
| 2 | Q9P2D6 | FAM135A | Protein FAM135A | ITKQPSSTSYNFTSSI | 8 |
| 2 | Q8WUM4 | PDCD6IP | Programmed cell dea | SYPFPQPPQQSYYPQQ | 8 |
| 2 | Q15735 | INPP5J | Phosphatidylinositol | SYRSHMEYTVSDHKPV | 8 |
| 2 | Q9BYW2 | SETD2 | Histone-lysine N-met | VYGVTSPYSQTTPPIV | 7 |
| 2 | Q86WV1 | SKAP1 | Src kinase-associated | LTSQDRRSYEFTATSP | 6 |
| 2 | Q86T96 | RNF180 | E3 ubiquitin-protein l | NSYSFQNPSSFDPSML | 6 |
| 3 | P10914 | IRF1 | Interferon regulatory | EVVPDSTSDLYNFQVS | 5 |
| 2 | Q9UQM7;Q13557;Q1: | CAMK2A;( | Calcium/calmodulin- | AYDFPSPEWDTVTPEA | 5 |
| 2 | Q96ST3 | SIN3A | Paired amphipathic h | QYSVTPSYQVSAMPQS | 5 |
| 2 | Q8IWB1 | ITPRIP | Inositol 1,4,5-trispho: | LISEYSLHVPSDQPTP | 4 |

**Table S2.** Radixin FERM ProP-PD results.

| Confidence | ProteinAcc | GeneName | ProteinName | Designed_peptide | Total NGS |
|---|---|---|---|---|---|
| 2 | Q6PJT7 | ZC3H14 | Zinc finger CCCH dom | YYDMESMVHADTRSFI | 5395 |
| 3 | Q6ZVL6 | KIAA1549L | UPF0606 protein KIA. | YYDFPAVETSKGLTER | 5005 |
| 2 | Q6S9Z5 | ZNF474 | Zinc finger protein 47 | SSSGSYSLQATNEAAF | 4034 |
| 2 | Q96BD5 | PHF21A | PHD finger protein 21 | TFTFPAPVQPVSLPSP | 3444 |
| 3 | P57082 | TBX4 | T-box transcription fa | SYSVQTMETVPYQPFP | 3347 |
| 2 | Q9HC84 | MUC5B | Mucin-5B | TWTVPAQTTTPMSTMS | 2862 |
| 3 | Q71RC2 | LARP4 | La-related protein 4 | SYQIYDVSGESNSAVS | 1967 |
| 2 | Q8TF21 | ANKRD24 | Ankyrin repeat doma | LSLLENERENTSYDVT | 1679 |
| 2 | P35398 | RORA | Nuclear receptor ROI | TYNISANGLTELHDDL | 1652 |
| 2 | Q2LD37 | KIAA1109 | Transmembrane prot | EWTLDQPVSQTRTTAI | 1610 |
| 2 | Q7Z570 | ZNF804A | Zinc finger protein 80 | YTLIHSEENTKDATTV | 1370 |
| 2 | Q14938 | NFIX | Nuclear factor 1 X-ty | NFSLADLESPSYYNIN | 1196 |
| 2 | Q6PCD5 | RFWD3 | E3 ubiquitin-protei | YFQVSRTQPDLPATTY | 1024 |
| 2 | Q8WZ42 | TTN | | VYSFEVQNPVGKDSCT | 1003 |
| 3 | Q86UR5 | RIMS1 | Regulating synaptic r | MYTLEHNDGSQSDTAV | 799 |
| 2 | P58107 | EPPK1 | Epiplakin {ECO:0000З | FFDPNTHENLTYVQLL | 741 |
| 2 | O60732 | MAGEC1 | Melanoma-associate | PVSSFFSYTLASLLQS | 591 |
| 3 | O75603 | GCM2 | Chorion-specific tran | SYELANPGYTNSSPYP | 536 |
| 2 | P01100 | FOS | Proto-oncogene c-Fo | AFTLPLLNDPEPKPSV | 460 |
| 2 | Q9P203 | BTBD7 | BTB/POZ domain-co | QEYPDFYDFSNAACRP | 449 |
| 2 | P20930 | FLG | Filaggrin | LYQVSTHEQSESTHGQ | 446 |
| 2 | Q9HBV1 | POPDC3 | Popeye domain-conta | FYQMSTPEIRRSPLTQ | 434 |
| 3 | Q96BF6 | NACC2 | Nucleus accumbens- | SYAVQEKPEPVPLESR | 417 |
| 2 | Q9Y264 | ANGPT4 | Angiopoietin-4 | SYTFLLPKSEPCPPGP | 415 |
| 2 | Q5TAQ9 | DCAF8 | DDB1- and CUL4-ass | EDTGHYSINDENRVHD | 383 |
| 2 | A8K2U0 | A2ML1 | Alpha-2-macroglobul | KYSMVELQDPNSNRIA | 368 |
| 2 | Q8N0W7 | FMR1NB | Fragile X mental reta | TYELAATESNPESSHP | 338 |
| 2 | Q9NRA8 | EIF4ENIF1 | Eukaryotic translatio | TPQNVPSRSGLPHMHS | 325 |
| 2 | Q99543 | DNAJC2 | DnaJ homolog subfar | FTPWTTEEQKLLEQAL | 295 |
| 2 | Q70E73 | RAPH1 | Ras-associated and p | SYSLDDVTAQLEQASL | 278 |
| 2 | Q8IUX7 | AEBP1 | Adipocyte enhancer-l | FPFTTVETYTVNFGDF | 259 |
| 2 | Q969S3 | ZNF622 | Zinc finger protein 62 | LEFADFYDFRSSYPDH | 258 |
| 2 | Q9Y4K1 | CRYBG1 | Beta/gamma crystall | SFVLPVESTQDVSSQV | 255 |
| 3 | Q86SP6 | GPR149 | Probable G-protein c | SYSLFLPTSNPDGDIN | 253 |
| 2 | P08631 | HCK | Tyrosine-protein kina | RDSETTKGSYSLSVRD | 247 |
| 3 | Q9HBZ2 | ARNT2 | Aryl hydrocarbon rec | SSYDLSQVPVPNLPAG | 224 |
| 3 | Q9H9V9 | JMJD4 | JmjC domain-contain | YDVTSPALCDTHLHPR | 221 |
| 2 | P20930 | FLG | Filaggrin | LYQVSTHEQSESAHGR | 217 |
| 3 | Q9Y5V3 | MAGED1 | Melanoma-associate | TYNFSQSLNANDLANS | 212 |
| 2 | Q7Z333 | SETX | Probable helicase se | AYELSQRSLDYVAQLR | 204 |
| 3 | Q02505 | MUC3A | Mucin-3A {ECO:0000 | TSSITSTYTVTSMTTT | 165 |
| 3 | P24928 | POLR2A | DNA-directed RNA p | SYSPTSPSYSPTSPSY | 161 |
| 2 | P14384 | CPM | Carboxypeptidase M | YYSIGRENYNQYDLNR | 154 |
| 3 | P78524 | ST5 | Suppression of tumo | SYQFPKLDRPTKQMRE | 148 |
| 2 | Q92797 | SYMPK | Symplekin | SFTPHQQAHIPNSIMT | 148 |
| 3 | Q7Z7G0 | ABI3BP | Target of Nesh-SH3 | TYDVFSSPTTSDEPEI | 144 |
| 3 | Q8IVT2 | MISP | Mitotic interactor an | ITGSYSVSESPFFSPI | 138 |
| 2 | O95359 | TACC2 | Transforming acidic | PTEELDYRNSYEIEYM | 124 |
| 3 | Q15032 | R3HDM1 | R3H domain-containi | YYSVIPPGQQNNLSSS | 122 |
| 2 | P60606 | CTXN1 | Cortexin-1 | MSATWTLSPEPLPPST | 120 |
| 2 | A7KAX9 | ARHGAP32 | Rho GTPase-activatir | DYHVTQLQPYFENGRV | 114 |
| 3 | Q8IZU9 | KIRREL3 | Kin of IRRE-like prote | YYSVNTFKEHHSTPTI | 113 |
| 2 | Q9NRA1 | PDGFC | Platelet-derived grov | KYDFVEVEEPSDGTIL | 101 |
| 2 | O60704 | TPST2 | Protein-tyrosine sulfo | YFQVNQNSTSSHLGSS | 97 |
| 2 | Q9BWV1 | BOC | Brother of CDO | LYTLPDDSTHQLLQPH | 84 |
| 2 | Q14643 | ITPR1 | Inositol 1,4,5-trispho | VYSLVSVPEGNDISSI | 84 |
| 2 | Q16665 | HIF1A | Hypoxia-inducible fac | FYELAHQLPLPHNVSS | 81 |
| 3 | O00167 | EYA2 | Eyes absent homolog | TYVLQEASHNVPNQSS | 80 |
| 2 | Q99569 | PKP4 | Plakophilin-4 | QYRVQECNYNRLQHAV | 77 |
| 2 | Q15149 | PLEC | Plectin | FFDPNTHENLTYRQLL | 72 |
| 2 | P55290 | CDH13 | Cadherin-13 | NYNLPIMVTDSGKPPM | 70 |
| 2 | Q6JVE5 | LCN12 | Epididymal-specific l | SYVLIPAAQPGQFTVD | 69 |
| 2 | Q2KHR3 | QSER1 | Glutamine and serine | TFCPPPLPKPSSTTPT | 66 |
| 2 | O95835 | LATS1 | Serine/threonine-pro | LYNISVPGLQTNWPQS | 65 |
| 2 | Q9UDY4 | DNAJB4 | DnaJ homolog subfar | AYEVLSDPKKREIYDQ | 58 |
| 2 | Q6N021 | TET2 | Methylcytosine dioxy | KSQMYQVEMNQGQSQG | 56 |
| 2 | Q96TA0 | PCDHB18P | Putative protocadher | SYTLFVRENNSPALHI | 56 |
| 2 | Q99700 | ATXN2 | Ataxin-2 | AYSPQQFPNQPLVQHV | 55 |
| 3 | Q3SXY7 | LRIT3 | Leucine-rich repeat, i | YTVIQESPEEGVRWSI | 53 |
| 3 | Q9H4A3 | WNK1 | Serine/threonine-pro | QYPVSQIPISTPHVST | 51 |
| 2 | Q03164 | KMT2A | Histone-lysine N-met | LGQNTSTSSNLQRTVV | 50 |
| 2 | O95835 | LATS1 | Serine/threonine-pro | THHKALQEIRNSLLPF | 50 |
| 2 | P22736 | NR4A1 | Nuclear receptor sub | FFSFSPPTGPSPSLAQ | 49 |
| 2 | Q86UW9 | DTX2 | Probable E3 ubiquitir | YTVNYTTHTQTNKTSS | 45 |
| 2 | Q9UPW0 | FOXJ3 | Forkhead box protein | SYTPVTSHPESVSQSL | 45 |
| 3 | P49005 | POLD2 | DNA polymerase delt | FDPTNYTLPQQPLHPC | 42 |
| 3 | P54296 | MYOM2 | Myomesin-2 | TYSVSVSDTDGVSSSF | 42 |
| 2 | P56181 | NDUFV3 | NADH dehydrogenase | TYTFLDLNLELSKFRM | 36 |
| 2 | P46939 | UTRN | Utrophin | TVTEVDMDLDSYQIAL | 34 |
| 3 | Q6EBC2 | IL31 | Interleukin-31 | LVSQNYTLPCLSPDAQ | 33 |
| 2 | Q07157 | TJP1 | Tight junction proteir | KYQINNISTVPKAIPV | 33 |
| 2 | Q9NU19 | TBC1D22B | TBC1 domain family | YFGFIEQYYDSRNEEH | 31 |
| 2 | Q16621 | NFE2 | Transcription factor I | NYTLPAAETPLALEPS | 30 |
| 2 | Q96BA8 | CREB3L1 | Cyclic AMP-responsiv | SYSLSGDSAPQSPLVP | 28 |
| 2 | Q9NS15 | LTBP3 | Latent-transforming | DTLPKQPCGSNPLPGL | 27 |
| 2 | Q9HAU6 | TPT1P8 | Putative translationa | TEDEVTESTIITSVDI | 27 |
| 2 | Q9Y5G0 | PCDHGB5 | Protocadherin gamm | QFTLQHVPDYRQNVYI | 26 |
| 3 | Q99614 | TTC1 | Tetratricopeptide rep | QDSSTGSYSINFVQNP | 25 |
| 2 | Q9H4I2 | ZHX3 | Zinc fingers and hom | SFSPSSKVPEVTCIPT | 25 |
| 2 | Q96L91 | EP400 | E1A-binding protein p | SYQIQQLMNRSPATGQ | 24 |
| 2 | Q92858 | ATOH1 | Protein atonal homol | QYLLHSPELGASEAAA | 23 |
| 2 | Q5H9K5 | ZMAT1 | Zinc finger matrin-ty | SLNQQENNSGSYSVES | 22 |
| 2 | Q9UH90 | FBXO40 | F-box only protein 40 | TYNFEPEQFSSGTVLA | 19 |
| 2 | Q8WUM4 | PDCD6IP | Programmed cell dea | SYPFPQPPQQSYYPQQ | 18 |
| 3 | Q9Y581 | INSL6 | Insulin-like peptide II | YSPYQFESPQTASPAR | 17 |
| 2 | Q8N2C7 | UNC80 | Protein unc-80 homo | NFTLPSPVLGMPSVPM | 17 |
| 2 | Q9P1A6 | DLGAP2 | Disks large-associate | QYSWSPTQHFNEERYS | 17 |
| 3 | P10914 | IRF1 | Interferon regulatory | LYNFQVSPMPSTSEAT | 16 |
| 2 | Q96ST3 | SIN3A | Paired amphipathic h | QYSVTPSYQVSAMPQS | 11 |
| 2 | P29074 | PTPN4 | Tyrosine-protein pho | NLSGYLSDYSFIPNQP | 9 |
| 2 | Q9Y698 | CACNG2 | Voltage-dependent c | MYTLSRDPLKAATTPT | 8 |
| 2 | Q8N813 | C3orf56 | Putative uncharacter | TYSLASPTLGGATSSH | 7 |
| 2 | Q8IZT6 | ASPM | Abnormal spindle-lik | YYSFIKQNNPKFSAVQ | 7 |
| 2 | Q6UW60 | PCSK4 | Proprotein convertas | PDLWANYDPLASYDFN | 5 |
| 2 | Q86US8 | SMG6 | Telomerase-binding | YYKFQNSDNPYYYPRT | 5 |
| 2 | A6NNM8 | TTLL13P | Tubulin polyglutamyl | PERVASDSWTECTLPS | 2 |

### **Table S3.** Moesin FERM domain ProP-PD results.

| Confidence | uniprot_accession | gene | protein_name | designed_peptide | Total NGS |
|---|---|---|---|---|---|
| 2 | Q96BD5 | PHF21A | PHD finger protein | TFTFPAPVQPVSLPSP | 16360 |
| 2 | P35398 | RORA | Nuclear receptor R | TYNISANGLTELHDDL | 11543 |
| 2 | Q9HC84 | MUC5B | Mucin-5B | TWTVPAQTTTPMSTMS | 9881 |
| 2 | Q7Z570 | ZNF804A | Zinc finger protein | YTLIHSEENTKDATTV | 9248 |
| 3 | Q8WZ42 | TTN | Titin | VYSFEVQNPVGKDSCT | 6854 |
| 3 | P57082 | TBX4 | T-box transcription | SYSVQTMETVPYQPFP | 4581 |
| 2 | Q2LD37 | KIAA1109 | Transmembrane pro | EWTLDQPVSQTRTTAI | 3525 |
| 2 | Q16665 | HIF1A | Hypoxia-inducible | FYELAHQLPLPHNVSS | 2263 |
| 2 | P58107 | EPPK1 | Epiplakin {ECO:000 | FFDPNTHENLTYVQLL | 1651 |
| 3 | Q8IZU9 | KIRREL3 | Kin of IRRE-like prot | YYSVNTFKEHHSTPTI | 1283 |
| 2 | Q6PJT7 | ZC3H14 | Zinc finger CCCH do | YYDMESMVHADTRSFI | 1196 |
| 2 | P01100 | FOS | Proto-oncogene c-F | AFTLPLLNDPEPKPSV | 1071 |
| 2 | Q92797 | SYMPK | Symplekin | SFTPHQQAHIPNSIMT | 1060 |
| 3 | O75385 | ULK1 | Serine/threonine-p | SFDFPKTPSSQNLLAL | 920 |
| 2 | Q8N0W7 | FMR1NB | Fragile X mental ret | TYELAATESNPESSHP | 917 |
| 2 | Q96BF6 | NACC2 | Nucleus accumbens | SYAVQEKPEPVPLESR | 825 |
| 3 | Q9Y5V3 | MAGED1 | Melanoma-associat | TYNFSQSLNANDLANS | 709 |
| 2 | Q86UR5 | RIMS1 | Regulating synaptic | MYTLEHNDGSQSDTAV | 671 |
| 3 | P20930 | FLG | Filaggrin | LYQVSTHEQSESAHGR | 632 |
| 2 | Q9HBV1 | POPDC3 | Popeye domain-cor | FYQMSTPEIRRSPLTQ | 585 |
| 2 | Q6PCD5 | RFWD3 | E3 ubiquitin-protei | YFQVSRTQPDLPATTY | 584 |
| 2 | A8K2U0 | A2ML1 | Alpha-2-macroglob | KYSMVELQDPNSNRIA | 556 |
| 2 | Q9P203 | BTBD7 | BTB/POZ domain-co | QEYPDFYDFSNAACRP | 513 |
| 2 | O75603 | GCM2 | Chorion-specific tra | SYELANPGYTNSSPYP | 445 |
| 3 | P78524 | ST5 | Suppression of tum | SYQFPKLDRPTKQMRE | 442 |
| 3 | Q9HBZ2 | ARNT2 | Aryl hydrocarbon re | SSYDLSQVPVPNLPAG | 418 |
| 2 | Q9H4A3 | WNK1 | Serine/threonine-pr | QYPVSQIPISTPHVST | 360 |
| 2 | Q9BWV1 | BOC | Brother of CDO | LYTLPDDSTHQLLQPH | 351 |
| 2 | Q9Y4K1 | CRYBG1 | Beta/gamma crystal | SFVLPVESTQDVSSQV | 349 |
| 2 | Q13237 | PRKG2 | cGMP-dependent p | TYDLNKPPEFSFEKAR | 344 |
| 3 | Q969S3 | ZNF622 | Zinc finger protein | LEFADFYDFRSSYPDH | 341 |
| 3 | Q9H9V9 | JMJD4 | JmjC domain-conta | YDVTSPALCDTHLHPR | 324 |
| 3 | Q96FA7 | ZBED6CL | ZBED6 C-terminal-li | VSEYSLNSPSPLQSPR | 300 |
| 2 | O60704 | TPST2 | Protein-tyrosine su | YFQVNQNSTSSHLGSS | 298 |
| 2 | Q99543 | DNAJC2 | DnaJ homolog subfa | FTPWTTEEQKLLEQAL | 297 |
| 3 | Q6S9Z5 | ZNF474 | Zinc finger protein | SSSGSYSLQATNEAAF | 296 |
| 2 | Q5SQI0 | ATAT1 | Alpha-tubulin N-ace | QYMLQKERVEPHQLAI | 294 |
| 2 | Q86UW9 | DTX2 | Probable E3 ubiqui | YTVNYTTHTQTNKTSS | 276 |
| 3 | Q15032 | R3HDM1 | R3H domain-contai | YYSVIPPGQQNNLSSS | 267 |
| 2 | P13611 | VCAN | Versican core prote | YLHITEPPSLSPDTKL | 260 |
| 3 | Q3SXY7 | LRIT3 | Leucine-rich repeat | YTVIQESPEEGVRWSI | 257 |
| 3 | Q8IVT2 | MISP | Mitotic interactor a | ITGSYSVSESPFFSPI | 231 |
| 2 | Q2KHR3 | QSER1 | Glutamine and serir | TFCPPPLPKPSSTTPT | 230 |
| 3 | Q86T96 | RNF180 | E3 ubiquitin-protei | NSYSFQNPSSFDPSML | 222 |
| 3 | P24928 | POLR2A | DNA-directed RNA p | SYSPTSPSYSPTSPSY | 200 |
| 3 | Q8NEY8 | PPHLN1 | Periphilin-1 {ECO:0 | YSPERSKSYSFHQSQH | 190 |
| 4 | P01243 | CSH1 | Chorionic somatom | YSFLHDSQTSFCFSDS | 177 |
| 2 | Q8N895 | ZNF366 | Zinc finger protein | CYEVEPYSPGLAPQSQ | 157 |
| 2 | Q02962 | PAX2 | Paired box protein | SNDPVGSYSINGILGI | 155 |
| 2 | Q9H4I2 | ZHX3 | Zinc fingers and hor | SFSPSSKVPEVTCIPT | 151 |
| 2 | Q8NEZ4 | KMT2C | Histone-lysine N-me | SYEVSSAPDVPSMGLV | 151 |
| 2 | Q70E73 | RAPH1 | Ras-associated and | SYSLDDVTAQLEQASL | 148 |
| 2 | Q8IZM8 | ZNF654 | Zinc finger protein | DYDLNQETSVIHKING | 147 |
| 3 | P49005 | POLD2 | DNA polymerase de | FDPTNYTLPQQPLHPC | 134 |
| 2 | P30048 | PRDX3 | Thioredoxin-depen | NWTPDSPTIKPSPAAS | 133 |
| 3 | Q8WUM4 | PDCD6IP | Programmed cell de | SYPFPQPPQQSYYPQQ | 130 |
| 3 | Q9Y581 | INSL6 | Insulin-like peptide | YSPYQFESPQTASPAR | 123 |
| 2 | Q8TAT6 | NPLOC4 | Nuclear protein loc | TFSISQNPFPIENRDV | 113 |
| 2 | Q99569 | PKP4 | Plakophilin-4 | QYRVQECNYNRLQHAV | 104 |
| 2 | Q53SZ7 | PRR30 | Proline-rich protein | SFSPSQPQNSSLPHSP | 101 |
| 2 | Q96T58 | SPEN | Msx2-interacting pr | TYSIRPEALHSPRAPL | 95 |
| 2 | Q07157 | TJP1 | Tight junction prot | KYQINNISTVPKAIPV | 90 |
| 2 | Q7Z333 | SETX | Probable helicase se | AYELSQRSLDYVAQLR | 89 |
| 2 | Q96K76 | USP47 | Ubiquitin carboxyl- | YSFNDQHVSRITQEDI | 84 |
| 3 | P10914 | IRF1 | Interferon regulato | LYNFQVSPMPSTSEAT | 79 |
| 3 | Q96BA8 | CREB3L1 | Cyclic AMP-respons | SYSLSGDSAPQSPLVP | 77 |
| 2 | Q6P0N0 | MIS18BP1 | Mis18-binding prot | YELPTLNQEQENFLAV | 75 |
| 3 | A7KAX9 | ARHGAP32 | Rho GTPase-activat | DYHVTQLQPYFENGRV | 74 |
| 3 | P30260 | CDC27 | Cell division cycle p | YSLNTDSSVSYIDSAV | 74 |
| 2 | O95835 | LATS1 | Serine/threonine-p | LYNISVPGLQTNWPQS | 74 |
| 2 | Q6JVE5 | LCN12 | Epididymal-specific | SYVLIPAAQPGQFTVD | 69 |
| 2 | Q5VTH2 | CFAP126 | Protein Flattop {EC | NWSPTKPTKESISSHE | 68 |
| 2 | Q9BVV6 | KIAA0586 | Protein TALPID3 | QYLFSPSREMPTFSGT | 67 |
| 2 | Q8WVN6 | SECTM1 | Secreted and transr | LWTPDSEPTPRPLALV | 66 |
| 3 | Q658P3 | STEAP3 | Metalloreductase S | KFTLPTDHALAEKTSH | 64 |
| 2 | O95835 | LATS1 | Serine/threonine-p | THHKALQEIRNSLLPF | 63 |
| 2 | O00443 | PIK3C2A | Phosphatidylinosito | YDLMVFPESDSQKRAL | 55 |
| 3 | Q9Y698 | CACNG2 | Voltage-dependent | MYTLSRDPLKAATTPT | 53 |
| 2 | P22736 | NR4A1 | Nuclear receptor su | FFSFSPPTGPSPSLAQ | 53 |
| 2 | Q96ST3 | SIN3A | Paired amphipathic | QYSVTPSYQVSAMPQS | 48 |
| 2 | Q01484 | ANK2 | Ankyrin-2 | EVSYEVTPKTTDVSTP | 47 |
| 2 | P29074 | PTPN4 | Tyrosine-protein ph | NLSGYLSDYSFIPNQP | 45 |
| 2 | Q96QE3 | ATAD5 | ATPase family AAA d | YSLNNDFVESSTSVLR | 42 |
| 3 | Q96I34 | PPP1R16A | Protein phosphatas | YQLSPLDSTTPHTLVH | 38 |
| 3 | Q5SVQ8 | ZBTB41 | Zinc finger and BTB | YTLTPQPANIVHPVRP | 34 |
| 2 | P54296 | MYOM2 | Myomesin-2 | TYSVSVSDTDGVSSSF | 34 |
| 2 | O60381 | HBP1 | HMG box-containin | QSDPTQSGMYQLSSDV | 32 |
| 3 | P06889 | IGLV3-1 | Immunoglobulin la | YELTQPPSVSVSPGQT | 31 |
| 2 | Q15735 | INPP5J | Phosphatidylinosito | SYRSHMEYTVSDHKPV | 31 |
| 2 | Q9NWT6 | HIF1AN | Hypoxia-inducible | DESQLRSYSFPTRPIP | 31 |
| 2 | Q9Y6N5 | SQOR | Sulfide:quinone oxi | SYEMLHVTPPMSPPDV | 31 |
| 2 | Q9UM54 | MYO6 | Unconventional my | YDFAPFLNNSPQQNPA | 31 |
| 2 | P20930 | FLG | Filaggrin | YTIQDEAYDTTDSLLE | 28 |
| 2 | Q6ZVL6 | KIAA1549L | UPF0606 protein K | YYDFPAVETSKGLTER | 26 |
| 2 | Q5VVP1;Q5TZJ5;Q5VYP | SPATA31A6;SP | Spermatogenesis-as | TYSLTGSTQQSRSLGA | 24 |
| 3 | Q13367 | AP3B2 | AP-3 complex subu | DYTFSRQPFSGDPHMV | 21 |
| 2 | P22105 | TNXB | Tenascin-X {ECO:000 | TGSSPDSLSLSWTVPQ | 21 |
| 2 | Q9Y264 | ANGPT4 | Angiopoietin-4 | SYTFLLPKSEPCPPGP | 21 |
| 2 | Q16236 | NFE2L2 | Nuclear factor eryth | EYSLQQTRDGNVFLVP | 20 |
| 2 | Q8TE57 | ADAMTS16 | A disintegrin and m | YDLVSAYEVDHRGDYV | 19 |
| 2 | P59025 | RTP1 | Receptor-transport | KLLEEEATTYTFSRAP | 18 |
| 2 | Q5H9K5 | ZMAT1 | Zinc finger matrin-t | SLNQQENNSGSYSVES | 18 |
| 2 | Q9NRJ4 | TULP4 | Tubby-related prote | LASQSSYSLLSPPDSA | 18 |
| 2 | O94898 | LRIG2 | Leucine-rich repeat | DYSITNTEELNLPADI | 17 |
| 2 | Q8N813 | C3orf56 | Putative uncharacte | TYSLASPTLGGATSSH | 17 |
| 2 | Q9ULJ8 | PPP1R9A | Neurabin-1 | NEYSVTGHYPLNLPSV | 16 |
| 2 | Q9NZM5 | NOP53 | Ribosome biogenes | SYNPSFEDHQTLLSAA | 16 |
| 2 | P01241 | GH1 | Somatotropin | YIPKEQKYSFLQNPQT | 15 |
| 2 | Q5VYJ5 | MALRD1 | MAM and LDL-recep | TESSVYSFSNPLYGTT | 15 |
| 2 | Q9UH90 | FBXO40 | F-box only protein | TYNFEPEQFSSGTVLA | 13 |
| 3 | Q5UIP0 | RIF1 | Telomere-associate | SKYAEYSFTSLPVPES | 12 |
| 2 | Q9UQM7;Q13557;Q13 | CAMK2A;CAM | Calcium/calmoduli | AYDFPSPEWDTVTPEA | 12 |
| 2 | A6NHM9 | MOXD2P | Putative DBH-like m | DSYDFNLQETRDLPSR | 12 |
| 2 | P10323 | ACR | Acrosin | HYDMETTELPELTSTS | 12 |
| 2 | Q9HD90 | NEUROD4 | Neurogenic differen | SPDLEKSYSFMPHYPS | 12 |
| 2 | Q0VF96 | CGNL1 | Cingulin-like protei | PVSYTFSKDSTVASQI | 11 |
| 2 | Q7Z6R9 | TFAP2D | Transcription facto | HHQSFHYEFQHSHPAV | 11 |
| 2 | P20930 | FLG | Filaggrin | YQVSSHEQSESTHGQT | 10 |
| 2 | Q9P2D6 | FAM135A | Protein FAM135A | ITKQPSSTSYNFTSSI | 10 |
| 2 | Q68DE3 | USF3 | Basic helix-loop-hel | YNLVSTSSMNTVACLP | 10 |
| 2 | O75427 | LRCH4 | Leucine-rich repeat | SYDLSDITQADLSRNR | 9 |
| 2 | Q86UW6 | N4BP2 | NEDD4-binding pro | PDDQVYSFLPSQDVNS | 9 |
| 2 | P10914 | IRF1 | Interferon regulato | EVVPDSTSDLYNFQVS | 8 |
| 2 | Q9H1K0 | RBSN | Rabenosyn-5 {ECO:0 | FYQLHSHYEEEHSGED | 8 |
| 2 | Q6ICB4 | PHETA2 | Sesquipedalian-2 {E | HYALSDSPADHMGFLR | 8 |
| 2 | Q86WV1 | SKAP1 | Src kinase-associate | LTSQDRRSYEFTATSP | 8 |
| 2 | Q9UQR1 | ZNF148 | Zinc finger protein | SLNFVTDNQTLPNQPA | 8 |
| 2 | Q92752 | TNR | Tenascin-R | TVPKDRTSYTLTDLEP | 8 |
| 4 | P42695 | NCAPD3 | Condensin-2 compl | SSYSLEQESNGEIEHV | 7 |
| 3 | Q8TDF6 | RASGRP4 | RAS guanyl-releasin | YTLSLEPETGCQLRHA | 7 |
| 2 | O15164 | TRIM24 | Transcription interr | GSYNLPSLPDIDCSST | 7 |
| 2 | Q9UPQ3 | AGAP1 | Arf-GAP with GTPas | YSVSSADQWSEATVIA | 7 |
| 2 | Q13950 | RUNX2 | Runt-related transc | SGSYQFPMVPGGDRSP | 6 |
| 2 | Q8WWN9 | IPCEF1 | Interactor protein f | SLSGTSYSFSSLENTV | 6 |
| 2 | P46937 | YAP1 | Transcriptional coa | SYSVPRTPDDFLNSVD | 6 |
| 2 | Q02505 | MUC3A | Mucin-3A {ECO:000 | TYSFSSSMSASSAGTT | 6 |
| 2 | Q04771 | ACVR1 | Activin receptor typ | YDVVPNDPSFEDMRKV | 6 |
| 2 | Q9ULC8 | ZDHHC8 | Probable palmitoyl | SSYSLQQASVLSEGPR | 6 |
| 2 | P01243 | CSH1 | Chorionic somatom | ETYIPKDQKYSFLHDS | 5 |
| 2 | O94952 | FBXO21 | F-box only protein | YNVEPQEISHPDVGRY | 5 |
| 2 | A6NCI8 | C2orf78 | Uncharacterized pr | YSVSSQPITETSVQVM | 5 |
| 2 | Q71RC2 | LARP4 | La-related protein 4 | GMILGPEDLSYQIYDV | 5 |
| 2 | Q9NS62 | THSD1 | Thrombospondin ty | KVYHVSQSPLTDTAID | 5 |

**Table S4.** Top enriched GO terms for the combined ERM ligand set.

| Category | Name | # in scree | # Proteome | Enrichment | P-value | Adj pval |
|---|---|---|---|---|---|---|
| Biological process | homophilic cell adhesion via plasma membrane adhesion mole | 15 | 111 | 11.04 | 4.35e-13 | 6.69e-10 |
| Biological process | calcium-dependent cell-cell adhesion via plasma membrane cel | 8 | 23 | 28.43.00 | 3.70e-12 | 2.85e-09 |
| Biological process | cell-cell adhesion via plasma-membrane adhesion molecules | 17 | 176 | 08.29 | 5.83e-12 | 2.99e-09 |
| Biological process | cell-cell adhesion | 22 | 371 | 05.25 | 1.96e-10 | 7.54e-08 |
| Biological process | cell adhesion | 21 | 425 | 04.04 | 1.38e-08 | 4.25e-06 |
| Biological process | nervous system development | 17 | 298 | 05.06 | 2.98e-08 | 7.64e-06 |
| Biological process | system development | 26 | 675 | 03.15 | 6.64e-08 | 1.43e-05 |
| Biological process | synapse organization | 11 | 131 | 07.26 | 7.43e-08 | 1.43e-05 |
| Biological process | cell adhesion | 28 | 775 | 03.35 | 8.72e-08 | 1.49e-05 |
| Biological process | biological adhesion | 28 | 780 | 03.33 | 1.00e-07 | 1.54e-05 |
| Molecular function | binding | 178 | 11557 | 01.26 | 6.00e-08 | 1.91e-05 |
| Molecular function | transcription regulatory region sequence-specific DNA binding | 24 | 676 | 2.9 | 9.01e-07 | 1.44e-04 |
| Molecular function | sequence-specific double-stranded DNA binding | 24 | 714 | 03.15 | 2.41e-06 | 1.50e-04 |
| Molecular function | calcium ion binding | 22 | 626 | 03.27 | 2.82e-06 | 1.50e-04 |
| Molecular function | RNA polymerase II transcription factor activity, sequence-specif | 31 | 1057 | 2.4 | 2.00e-06 | 1.50e-04 |
| Molecular function | protein binding | 111 | 6373 | 01.42 | 2.14e-06 | 1.50e-04 |
| Molecular function | double-stranded DNA binding | 25 | 793 | 02.58 | 4.92e-06 | 2.24e-04 |
| Biological process | interferon-gamma-mediated signaling pathway | 7 | 68 | 08.41 | 1.75e-06 | 2.32e-04 |
| Biological process | regulation of histone H3-K27 acetylation | 2 | 3 | 54.48.00 | 1.81e-06 | 2.32e-04 |
| Biological process | developmental process | 87 | 4597 | 01.55 | 2.00e-06 | 2.37e-04 |

**Table S5.** Merlin FERM domain ProP-PD results. Outcome of clonal phage ELISA assay are indicated, when available.

| Confidence | protein_accession | protein_gene | protein_name | protein_organism | library_peptide | Total NGS | Confirmed by phage ELISA |
|---|---|---|---|---|---|---|---|
| 2 | A6ND36 | FAM83G | Protein FAM83G | Homo sapiens (Human | AEPLPSLEYWPQKSD | 43881 | yes |
| 2 | O75603 | GCM2 | Chorion-specific tı | Homo sapiens (Human | SYELANPGYTNSSPYF | 3325 | NA |
| 2 | O95835 | LATS1 | Serine/threonine-p | Homo sapiens (Human | THHKALQEIRNSLLPF | 599 | NA |
| 2 | Q6ZVL6 | KIAA1549L | UPF0606 protein I | Homo sapiens (Human | YYDFPAVETSKGLTER | 31 | yes |
| 2 | Q96BD5 | PHF21A | PHD finger protein | Homo sapiens (Human | TFTFPAPVQPVSLPSP | 17 | NA |
| 2 | Q8IZU9 | KIRREL3 | Kin of IRRE-like prc | Homo sapiens (Human | YYSVNTFKEHHSTPTI | 7 | NA |
| 2 | Q9Y2H9 | MAST1 | Microtubule-assoc | Homo sapiens (Human | LFRKITKQSNLLHTSR | 6 | NA |
| 2 | Q7Z570 | ZNF804A | Zinc finger protein | Homo sapiens (Human | YTLIHSEENTKDATTV | 4 | NA |
| 1 | Q969S3 | ZNF622 | Zinc finger protein | Homo sapiens (Human | LEFADFYDFRSSYPDH | 57 | yes |
| 1 | Q9P2E9 | RRBP1 | Ribosome-binding | Homo sapiens (Human | RAATRLQELLKTTQEC | 24 | yes |
| 1 | Q9NZM5 | NOP53 | Ribosome biogene | Homo sapiens (Human | FYDLWASDNPLDRPI | 14 | yes |
| 1 | P10619 | CTSA | Lysosomal protect | Homo sapiens (Human | HYWFVESQKDPENSI | 3 | yes |
| 1 | Q9NQU5 | PAK6 | Serine/threonine-p | Homo sapiens (Human | RTWHAQISTSNLYLPC | 1 | yes |
| 1 | O60271 | SPAG9 | C-Jun-amino-term | Homo sapiens (Human | KKRSSIWQFFSRLFSS | 1 | yes/no |
| 1 | Q7Z6Z7 | HUWE1 | E3 ubiquitin-prote | Homo sapiens (Human | LTENQLQLSVEVLTSH | 1 | yes/no |
| 1 | P38398 | BRCA1 | Breast cancer type | Homo sapiens (Human | LQNTFKVSKRQSFAPI | 1 | no |
| 1 | Q9UGU0 | TCF20 | Transcription facto | Homo sapiens (Human | LPKNPPPKRATEMQS | 1 | no |
| 1 | Q5GLZ8 | HERC4 | Probable E3 ubiqu | Homo sapiens (Human | SHEINPRKVFELMGSI | 2 | no |
| 1 | Q8IVT5 | KSR1 | Kinase suppressor | Homo sapiens (Human | LTRLRRTESVPSDINN | 2 | no |
| 1 | Q5VU43 | PDE4DIP | Myomegalin | Homo sapiens (Human | KEQESIIQQLQTSLHD | 1 | no |
| 1 | Q9P267 | MBD5 | Methyl-CpG-bindi | Homo sapiens (Human | PVNQNPVIINPTSFHS | 1 | no |
| 1 | Q03164 | KMT2A | Histone-lysine N-m | Homo sapiens (Human | LSSLESSRRVHTSTPS | 20 | no |
| 1 | Q8NA75 | DCAF4L2 | DDB1- and CUL4-a | Homo sapiens (Human | LLTTIPSPYPASENDI | 1 | no |

**Table S6.** Detection of binding hotspot residues in F3a and F3b binding sites using computational alanine scanning.

| A | F3b binders | | | | | |
|---|---|---|---|---|---|---|
| | **MISP** | | **TBX4** | | **KIRREL** | |
| | DDG* | DG** | DDG | DG | DDG | DG |
| **M285A** | 0.85 | 0.59 | 1.40 | 0.42 | - | - |
| **H288A** | 2.17 | -0.06 | 3.14 | -0.14 | 2.17 | -0.01 |

| B | F3a binders | | | |
|---|---|---|---|---|
| | **ZNF622** | | **BTBD7** | |
| | DDG | DG | DDG | DG |
| **K211A** | 1.59 | -0.75 | 1.50 | 0.79 |
| **I238A** | 1.16 | 0.72 | - | - |
| **F267A** | 1.85 | 2.00 | 1.74 | 1.95 |

* DDG indicates the predicted change upon mutation to alanine in binding affinity

** DG indicates the predicted change upon mutation to alanine in stability.

**Table S7:** Binding of representative peptides to moesin mutants targeting the F3a (M285A/H288A) and F3b pockets (F267A/K211A).

| **Name** | **FITC-labeled probe** | **$K_D$ (µM)** | |
|---|---|---|---|
| | | **Moesin M285A/H288A** | **Moesin F267A/K211A** |
| **$LATS1_{73-88}$** | THHKALQEIRNSLLPF | 1.3 ± 0.1 | 5.3 ± 0.4 |
| **$EBP50_{343-358}$** | APQMDWSKKNELFSNL-coo- | 3.1 ± 0.2 | 12.6 ± 1.1 |
| **$ZNF622_{341-356}$** | LEFADFYDFRSSYPDH | 1.3 ± 0.1 | 5.0 ± 0.3 |
| **$KIRREL3_{637-652}$** | YYSVNTFKEHHSTPTI | 0.84 ± 0.05 | 0.14 ± 0.01 |
| **$TBX4_{428-443}$** | SYSVQTMETVPYQPFP | 6.9 ± 0.4 | 4.5 ± 0.2 |
| **$NOP53_{192-207}$** | FYDLWASDNPLDRPLV | 2.2 ± 0.1 | 27.3 ± 0.2 |

**Table S8.** Affinity determinations through FP competition experiments. The FITC-labeled probe peptides are indicated with names, and the competing peptides are indicated with names and peptide sequences.

| **Name** | **Peptide** | **FITC-EBP50 $K_D$' (μM)** | **FITC-ZNF622 $K_D$' (μM)** | **FITC-KIRREL3 $K_D$' (μM)** | **FITC-TBX4 $K_D$' (μM)** | **FITC-NOP53 $K_D$' (μM)** |
|---|---|---|---|---|---|---|
| EBP50$_{343-358}$ | APQMDWSKKNEL**F**SN**L**-COO- | **2.1 ± 0.1** | 3.5 ± 0.3 | 2.5 ± 0.1 | - | 3.1 ± 0.3 |
| **Motif: E[Y/F]xDFYDF** | | | | | | |
| ZNF622$_{341-356}$ | LEFAD**FYDF**RSSYPDH | 5.2 ± 0.1 | **1.03 ± 0.04** | 1.61 ± 0.06 | - | 4.13 ± 0.04 |
| BTBD7$_{940-950}$ | QEYPD**FYDF**SNAAARP | 2.6 ± 0.1 | 2.53 ± 0.04 | 1.34 ± 0.06 | - | 2.7 ±0.1 |
| **Motif: YxV** | | | | | | |
| KIRREL3$_{637-652}$ | Y**Y**S**V**NTFKEHHSTPTI | 14 ±1 | 52 ± 1 | **0.09 ± 0.01** | 0.15 ± 0.09 | 59.3 ±0.3 |
| TBX4$_{428-443}$ | S**Y**S**V**QTMETVPYQPFP | 176 ±10 | 148 ± 6 | 33 ± 1 | **1.0 ± 0.1** | 144 ± 5 |
| MISP$_{595-610}$ | ITGS**Y**S**V**SESPFFSPI | 34.4 ±0.5 | | 2.3 ± 0.1 | 1.8 ± 0.1 | |
| **Motif: FY[D/E]L(4-5x)PLxxx[L/V]** | | | | | | |
| NOP53$_{192-207}$ | **FYDL**WASDN**PL**DRP**L**V | N.D. | 197 ± 5 | 117 ± 3 | | **0.78 ± 0.05** |
| HIF1A$_{37-52}$ | **FYEL**AHQL**PL**PHN**V**SS | N.D. | 248 ± 7 | 430 ± 20 | | 3.0 ± 0.4 |

N.D: Not determinable.
