## Supplemental Figures 1-7 for "Short linear motif based interactions and dynamics of the ezrin, radixin, moesin and merlin FERM domains"

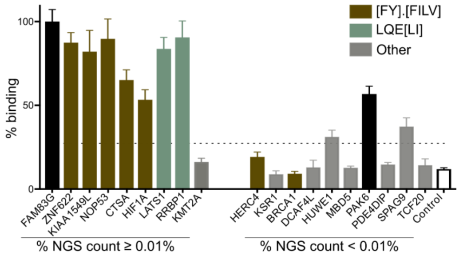


**Figure S1: Validation of putative merlin ligands through clonal phage ELISA.** Merlin ligands suggested by the phage display selection were tested for binding through clonal phage ELISA. The % binding was normalized to the result of FAM83G, which gave the highest signal. The negative control (control) indicates the result of a clonal phage ELISA against M13 phage displaying no peptides. HUWE1 and SPAG9 were considered borderline cases.

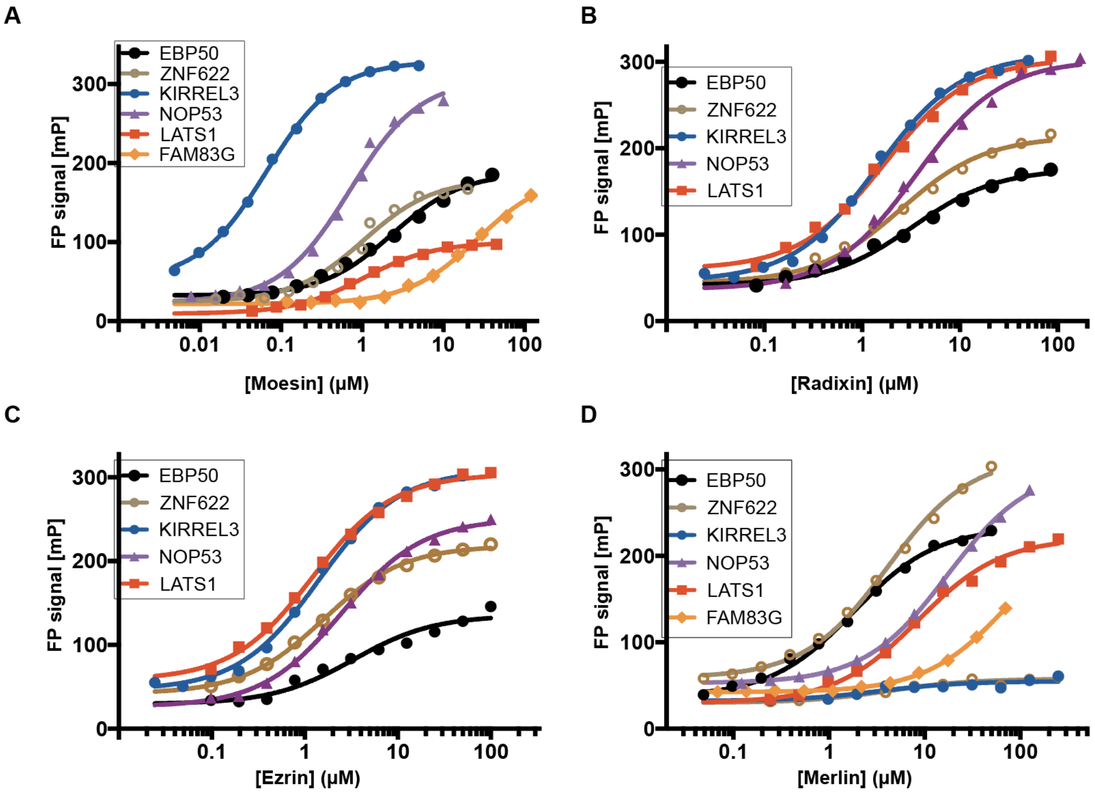


**Figure S2. Raw FP binding data of the normalized data shown in Fig 3A. (A-D)** FITC-labeled probe peptides (5 nM) were titrated with increasing concentrations of (**A**) moesin, (**B**) ezrin, (**C**) radixin, and (**D**) merlin FERM domains. The results were fitted to a quadratic equation for 1:1 binding (see Table 2 for $K_D$ values, n=3).

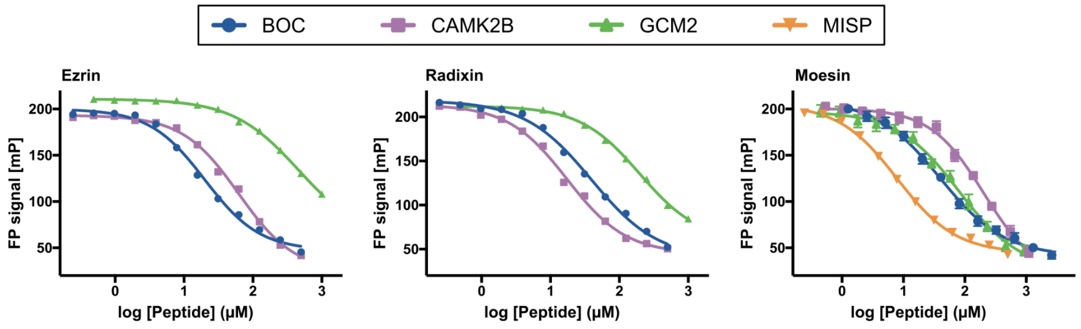


**Figure S3: Competition FP assay demonstrates binding of additional ligand set for FERM domains.** Kirrel3 was used as a probe in these competition FP experiments, while unlabeled peptides compete for binding to Ezrin, Radixin and Moesin FERM domains. (See Table 2 for $K_D$' values).

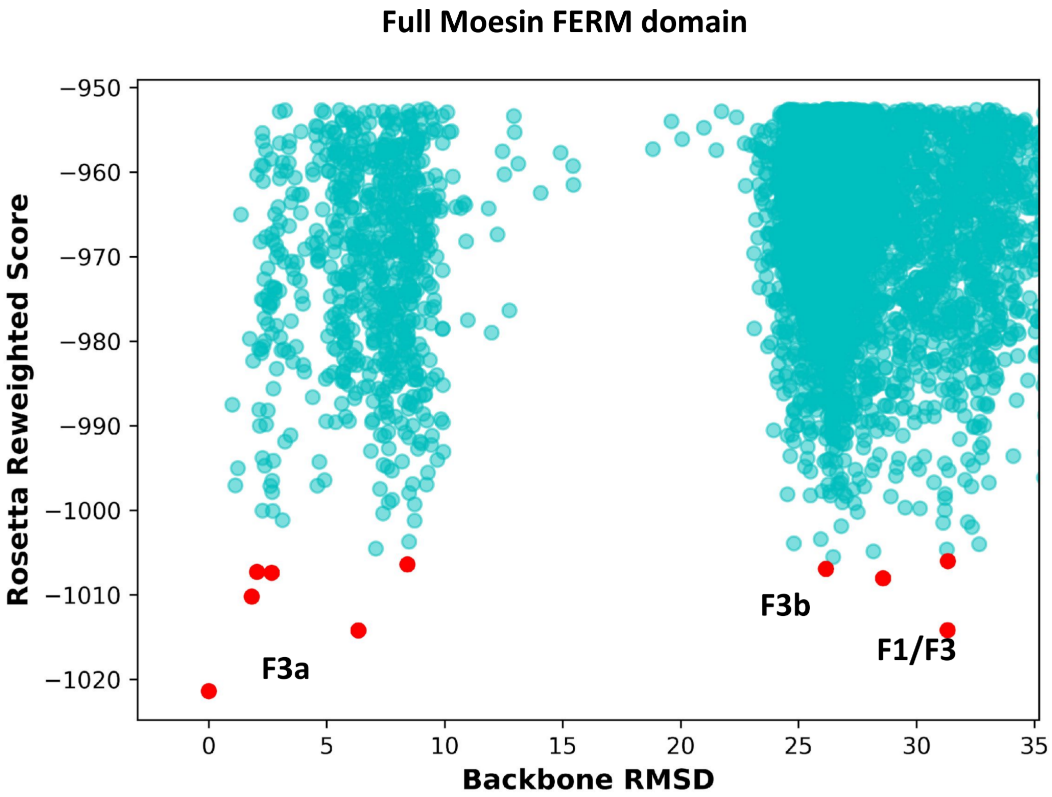


**Figure S4. ZNF622 simulation on full FERM domain.** Energy landscape of the global docking simulation of the ZNF622$_{341\text{-}356}$ peptide onto the full FERM domain shows clear distribution of the low energy models between two binding sites: left funnel - F3a site, right funnel - the F1/F3 cleft. The RMSD is calculated from the top-scoring model of the simulation. Cluster centers are shown with red dots.

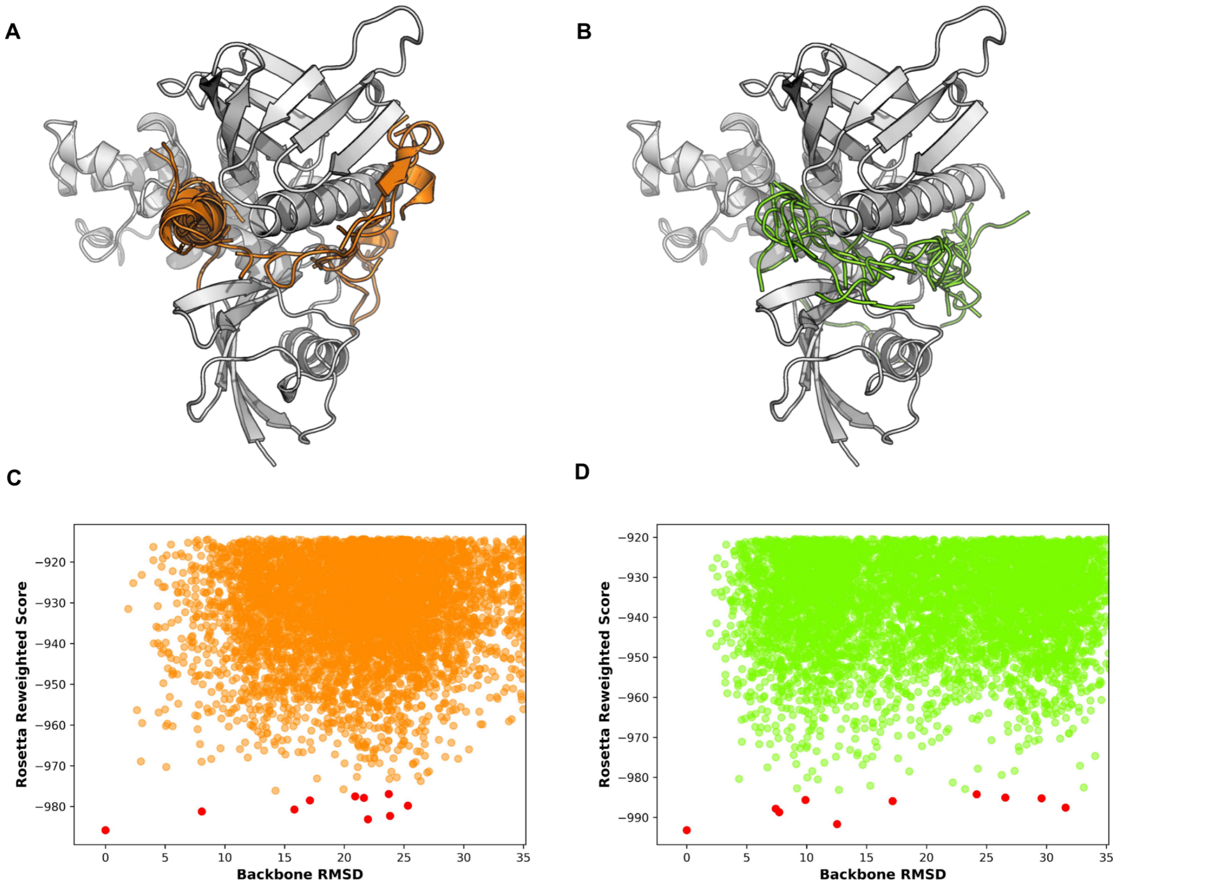


**Figure S5. NOP53$_{192-207}$ and HIF1A.** Top 20 models from global docking of **(A)** NOP53$_{192-207}$ and **(B)** HIF1a are shown to predominantly occupy the interdomain sites. **(C)-(D)**The energy landscapes of both NOP53$_{192-207}$ (orange) and HIF1a (green) simulations show a diffuse distribution of binding modes. The RMSD is calculated from the top-scoring model of the simulation. Cluster centers are shown with red dots.

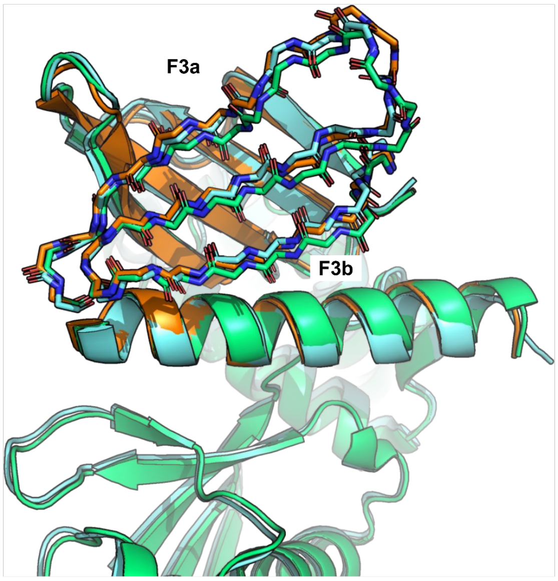


**Figure S6:** Opening of F3b pocket by relaxing the F3a bound structure in presence of a peptide bound to F3b (See Figure 7F). Green - F3a bound structure (1EF1), cyan - F3b bound structure (6TXS), orange - 1EF1 after Rosetta Relax simulation with a peptide superimposed on the F3b binding site.

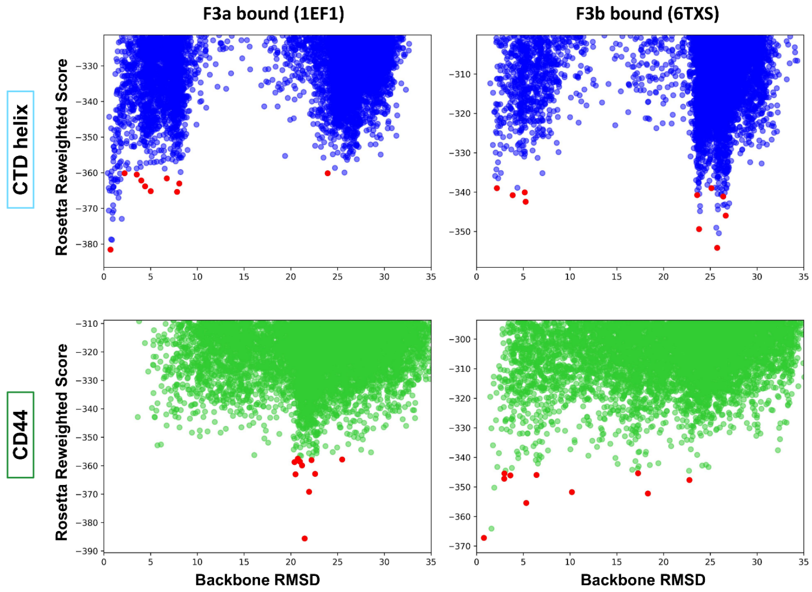


**Figure S7. Control runs with known ligands confirm the conformational change caused by the F3a binders.** The F3b binding site is closed by binding to F3a: Energy landscapes from PIPER-FlexPepDock global docking simulations on the F3 subdomain. Global docking of CD44 to moesin does not identify the F3b binding site when an F3a bound structure is used. However CTD helix (bound to F3a) successfully samples its binding site when F3b bound structure is used. Only the F3 subdomain was used in the simulations. Top 10 cluster representatives of each simulation are shown with red dots. The RMSD is calculated relative to the native site of the peptide (F3b for the CD44 and F3a for the CTD fragment).
